# Fidelity of parent-offspring transmission and the evolution of social behavior in structured populations

**DOI:** 10.1101/082503

**Authors:** F. Débarre

## Abstract

The theoretical investigation of how spatial structure affects the evolution of social behavior has mostly been done under the assumption that parent-offspring strategy transmission is perfect, *i.e.*, for genetically transmitted traits, that mutation is very weak or absent. Here, we investigate the evolution of social behavior in structured populations under arbitrary mutation probabilities. We consider populations of fixed size *N*, structured such that in the absence of selection, all individuals have the same probability of reproducing or dying (neutral reproductive values are the all same). Two types of individuals, *A* and *B*, corresponding to two types of social behavior, are competing; the fidelity of strategy transmission from parent to offspring is tuned by a parameter *μ*. Social interactions have a direct effect on individual fecundities. Under the assumption of small phenotypic differences (implying weak selection), we provide a formula for the expected frequency of type *A* individuals in the population, and deduce conditions for the long-term success of one strategy against another. We then illustrate our results with three common life-cycles (Wright-Fisher, Moran Birth-Death and Moran Death-Birth), and specific population structures (graph-structured populations). Qualitatively, we find that some life-cycles (Moran Birth-Death, Wright-Fisher) prevent the evolution of altruistic behavior, confirming previous results obtained with perfect strategy transmission. We also show that computing the expected frequency of altruists on a regular graph may require knowing more than just the graph’s size and degree.

## 1 Introduction

Most models on the evolution of social behavior in structured populations study the outcome of competition between individuals having different strategies and assume that strategy transmission from parents to their offspring is almost perfect (*i.e.*, when considering genetic transmission, that mutation is either vanishingly small or absent). This is for instance illustrated by the use of fixation probabilities to assess evolutionary success (*e.g.*, Rousset & Billiard, 2000; Rousset, 2003; Nowak et al., 2004; Nowak, 2006; Ohtsuki et al., 2006). Yet, mutation has been shown to affect the evolutionary fate of social behavior (Frank, 1997; Tarnita et al., 2009) and is, more generally, a potentially important evolutionary force. Here, we explore the role of imperfect strategy transmission—genetic or cultural—from parents to offspring on the evolution of social behavior, when two types of individuals, with different social strategies, are competing. We are interested in evaluating the long-term success of one strategy over another.

A population in which mutation is not close (or equal) to zero will spend a non-negligible time in mixed states (*i.e.*, in states where both types of individuals are present), so instead of fixation probabilities, we need to consider long-term frequencies to assess evolutionary success (Tarnita et al., 2009; Wakano & Lehmann, 2014; Tarnita & Taylor, 2014). We will say that a strategy is favored by selection when its expected frequency is larger than what it would be in the absence of selection.

Obviously, lowering the fidelity of parent-offspring strategy transmission—*e.g.*, by increasing the probability of mutation—reduces the relative role played by selection. But in a spatially structured population, the fidelity of parent-offspring strategy transmission also affects the spatial clustering of different strategies, and in particular whether individuals that interact with each other have the same strategy or not; this effect takes place even in the absence of selection. Consequently, the impact of imperfect strategy transmission may differ according to how the population is structured.

In this study, we consider populations such that, in the absence of selection (when social interactions have no effect on fitness), all individuals have equal chances of reproducing, and equal chances of dying. In other words, in such a population of size *N*, the neutral reproductive value of each site is 1/*N* (Taylor, 1990; Maciejewski, 2014; Tarnita & Taylor, 2014). We provide a formula that gives the long-term frequency of a social strategy in any such population, for arbitrary mutation rates, and for any life-cycle (provided population size remains equal to *N*). This formula is a function of the probabilities that pairs of individuals are identical by descent. These probabilities are obtained by solving a linear system of equations, and we present explicit solutions for population structures with a high level of symmetry (structures that we call “n-dimensional graphs”). We finally illustrate our results with widely used updating rules (Moran models, Wright-Fisher model) and specific population structures.

## 2 Models and Methods

### 2.1 Population structures

We consider a population of fixed size *N*, where each individual inhabits a site corresponding to the node of a graph 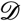; each site hosts exactly one individual. The edges of the graph, {*d*_*ij*_}_1≤*i*, *j*≤*N*_, define where individuals can send their offspring to. We consider graphs 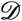 that are connected, *i.e.*, such that following the edges of the graph, we can go from any node to any other node (potentially via other nodes). This simply means that there are not completely isolated sub-populations. Another graph, *ℰ*, with the same nodes as graph 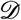 but with edges {*e*_*ij*_}_1≤*i*, *j*≤*N*_, defines the social interactions between the individuals; *ℰ* can be the same graph as 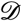, but does not have to be (Taylor et al., 2007a; Ohtsuki et al., 2007; Débarre et al., 2014). The edges of the two graphs can be weighted (*i.e.*, *d*_*ij*_ and *e*_*ij*_ can take any non-negative value) and directed (*i.e.*, we can have *d*_*ij*_ ≠ *d*_*ji*_ or *e*_*ij*_ ≠ *e*_*ji*_ for some sites *i* and *j*). For instance, dispersal in a subdivided population is represented by a weighted graph (the probability of sending offspring to a site in the same deme as the parent is different from the probability of sending offspring to a site in a different deme.) Finally, we denote by **D** and **E** the adjacency matrices of the dispersal and interaction graphs, respectively 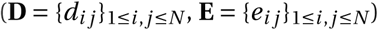.

#### Regular dispersal graphs

In this study, we focus on dispersal graphs that are regular, *i.e.*, such that for all sites *i*, the sum of the edges to *i* and the sum of the edges from *i* are both equal to *v*:

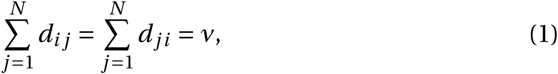

where *v* is called *degree* of the graph when the graph is unweighted. All the graphs depicted in the article (figures 1 and 3) satisfy eq. (1), and then are regular. Note that there is no specific constraint on the interaction graph *ℰ*.

**Figure 1:**
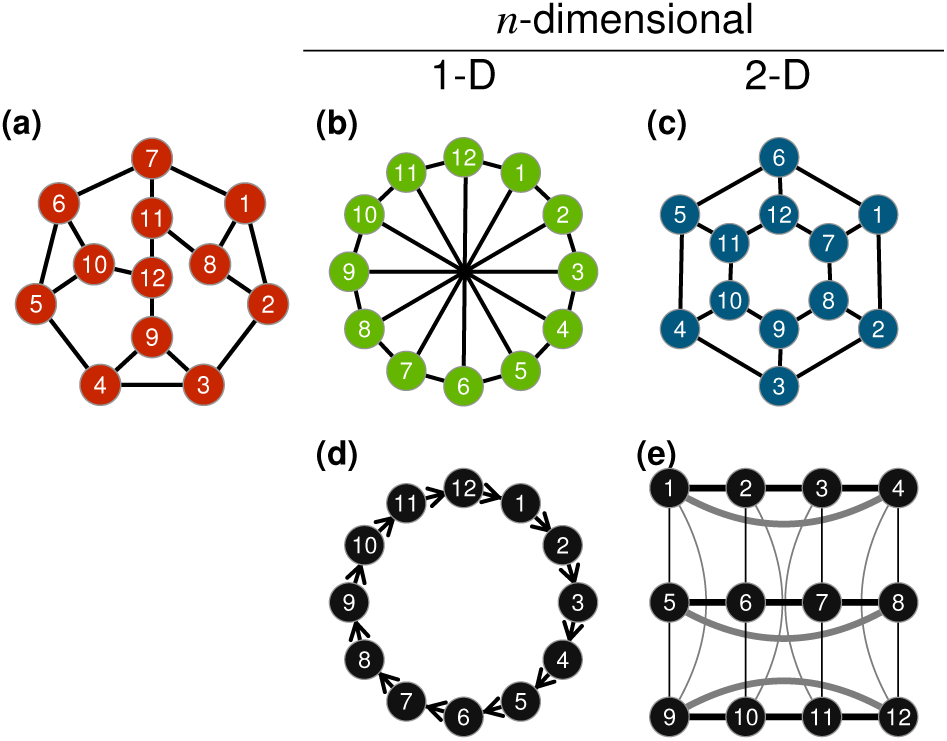
Examples of regular graphs of size 12. The graphs on the first line are unoriented and unweighted graphs of degree *v* = 3; Graph (d) is oriented, graph (e) is weighted. (a) is the Frucht graph, and has no symmetry. Graphs (b) and (d) are one-dimensional, graphs (c) and (e) are two-dimensional (see main text).

**Figure 2:**
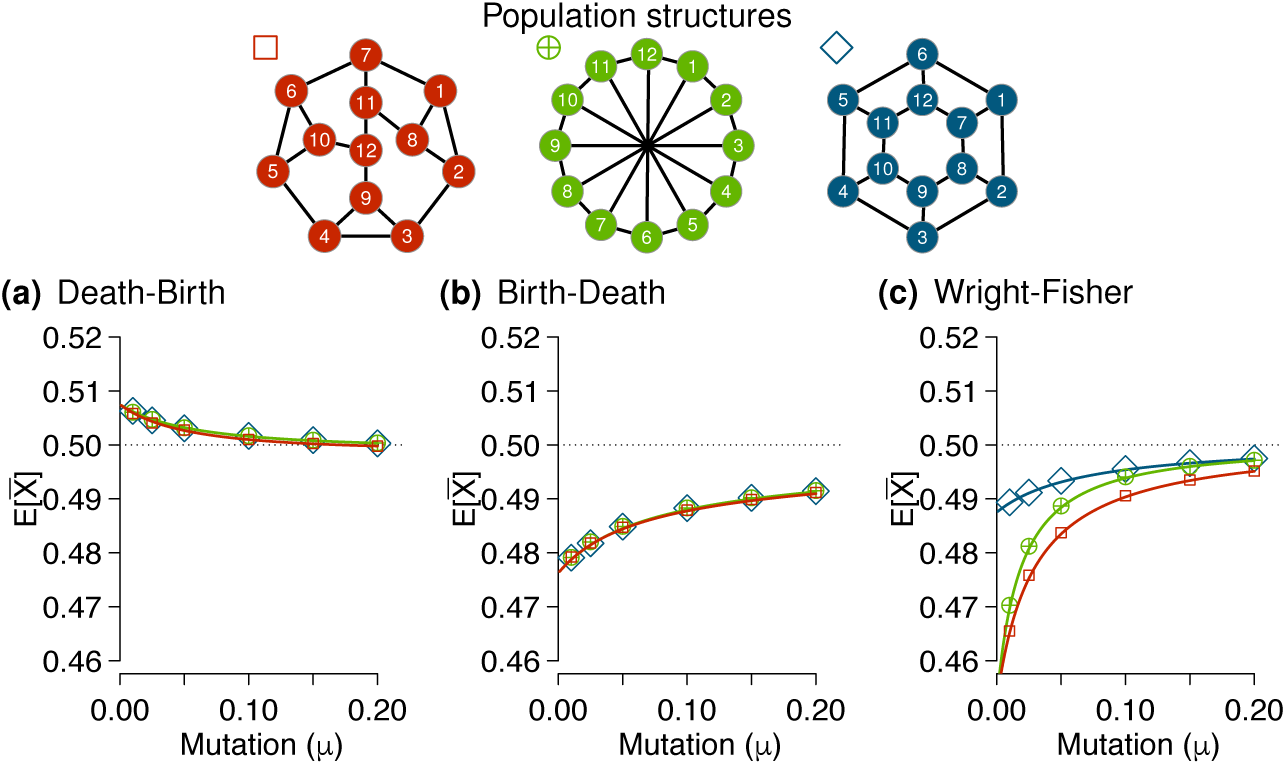
Expected frequency of type-A individuals 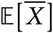, depending on population structure (legend on the first line), updating rule ((a): Moran Death-Birth, (b): Moran Birth-Death, (c): Wright-Fisher), and mutation probability *μ* (horizontal axis): Comparison between the theoretical prediction (curves) and the outcomes of numerical simulations (points). The horizontal dotted gray line corresponds to *p*, the expected frequency of type-*A* individuals when there is no selection (i.e.,when *δ* = 0). Other parameters: *δ*=0.005, *p* = 1/2, b = 8, c = 1.

**Figure 3:**
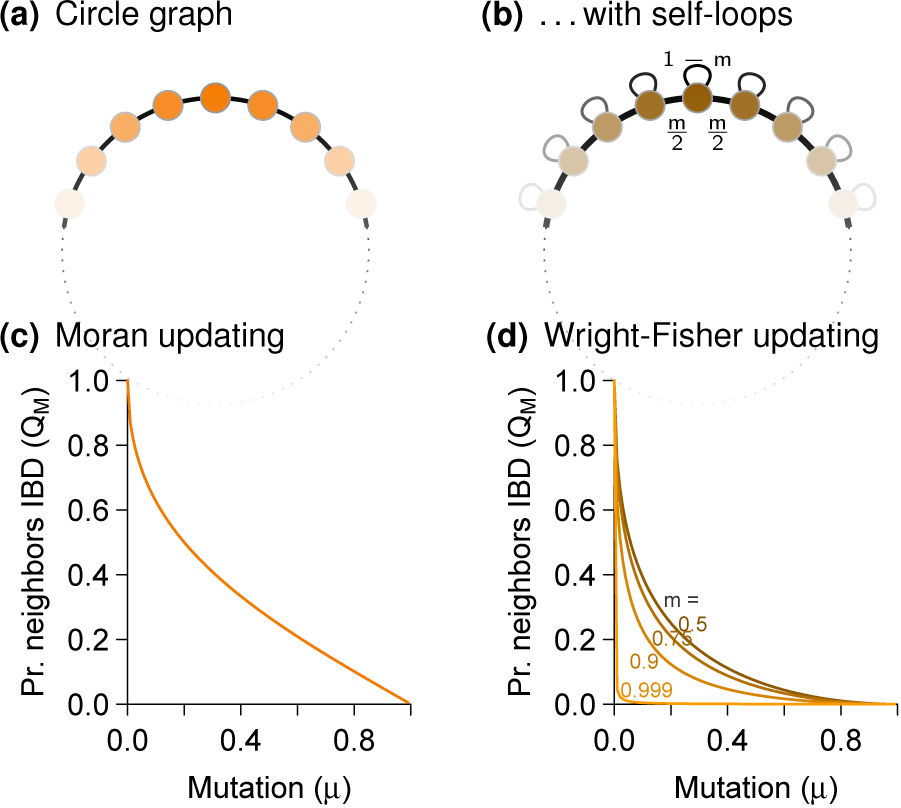
Circle graphs, without (a) or with self-loops ((b); the weight of the self-loop is 1 − *m*), and Probability that two neighbors on the graph are identical by descent, as function of the mutation probability *μ*, for the Moran updating on an infinite circle graph (c), and for the Wright-Fisher updating on an infinite circle graph with self loops (d). In (d), emigration probabilities *m* take values 0.5, 0.75, 0.9, 0.999 (increasingly lighter curves).

More detailed results are then obtained for regular graphs that display some level of symmetry, that we now describe:

#### Transitive dispersal graphs

A transitive graph is such that for any two nodes *i* and *j* of the graph, there is an isomorphism that maps *i* to *j* (Taylor et al., 2007a,b): the graph looks the same from every node. In other words, the dispersal graph is transitive when it is “homogeneous” (*sensu* Taylor et al., 2007a), *i.e.*, when all nodes have exactly the same properties in terms of dispersal. In figure 1, graphs (b)–(e) are transitive. On the other hand, all the nodes of graph (a) are different (for instance, node 9 is in a triangle while node 12 is not), so this regular graph is not transitive.

#### Transitive undirected dispersal graphs

A graph is undirected if for any two nodes *i* and *j*, the weight of the edge from *i* to *j* is equal to the weight of the edge from *j* to *i* (*i.e.*, there is no need to use arrows when drawing the edges of the graph). The dispersal graph is undirected when for all sites *i* and *j*, *d*_*ij*_ = *d*_*ji*_. In figure 1, graphs (b), (c), (e) are both transitive and undirected.

#### “*n*-dimensional” dispersal graphs

We call “*n*-dimensional graphs” transitive graphs whose nodes can be relabelled with *n*-long indices, such that the graph is unchanged by circular permutation of the indices in each dimension (see eq. (2)). We denote by 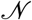 the ensemble of node indices: 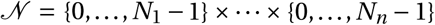, with 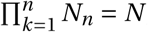; numbering is done modulo *N*_*k*_ in dimension *k*. Then for all indices *i*, *j* and *k* of *N*, node labeling is such that for all edges (modulo the size of each dimension),

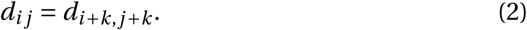

In figure 1, graphs (b) and (d) are 1-dimensional: we can label their nodes such that the adjacency matrices are circulant. Graphs (c) and (e) are 2-dimensional: the adjacency matrices are block-circulant, with each block being circulant. In 1(c), one dimension corresponds to the angular position of a node (*N*_1_ = 6 positions), and the other dimension to the radial position of a node (*N*_2_ = 2 positions, inner or outer hexagon). In 1(e), one dimension corresponds to the horizontal position of a node (*N*_1_ = 4 positions) and the other to the vertical position of a node (*N*_2_ = 3 positions). Condition eq. (2) may sound strong, but is satisfied for the regular population structures classically studied, like stepping-stones (*e.g.*, cycle graphs, lattices), or island models (Taylor, 2010; Taylor et al., 2011).

### 2.2 Types of individuals and social interactions

There are two types (*A* and *B*) of individuals in the population, corresponding to two strategies of social behavior. There are no mixed strategies: an individual of type *A* plays strategy *A*, and individuals do not change strategies. The indicator variable *X*_*i*_ represents the type of the individual present at site *i*: *X*_*i*_ is equal to 1 if the individual at site *i* is of type *A*, and *X*_*i*_ is equal to 0 otherwise 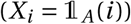. A *N*-long vector *X* gathers the identities of all individuals in the population, and 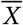 is the population average of 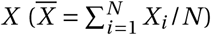. The ensemble of all possible states is 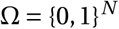.

Individuals in the population reproduce asexually. Fecundities are affected by social interactions, and are gathered in a *N*-long vector *f*. We assume that the genotype-phenotype map is such that the two types *A* and *B* are close in phenotype space: the individual living at site *i* expresses a phenotype *δX*_*i*_, with *δ* ≪ 1 (a feature called “*δ*-weak selection” by Wild & Traulsen (2007)).

An individual’s fecundity depends on the phenotypes of the individuals it interacts with and on its own phenotype (*δX*_*i*_ for the individual at site *i*). Without loss of generality, we can write the fecundity of the individual living at site *i* as

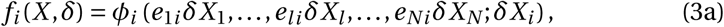

where the *N* first arguments correspond to the potential interactants (an individual can also be interacting with itself if *e*_*ii*_ > 0, which can occur for instance with a common-good), and the last (*N* + 1) argument is the phenotype of the focal individual.

Scaling fecundities such that the baseline in the absence of selection is equal to 1, a first-order expansion of eq. (3a) yields

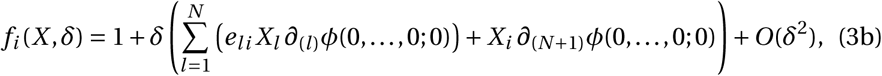

where *∂*_(*n*)_*ϕ_i_* represents the partial derivative of *ϕ_i_* with respect to its *n*^th^ element.

We do not need specify a particular shape for *ϕ_i_*; the only assumption that we make is that it does not matter where the interactions actually take place, only that they do take place, and that it does not matter either where the focal individual is (*i.e.*, there are no external sources of heterogeneity affecting individual fecundities). So for all *i* and *l*, 1 ≤ *i*, *l* ≤ *N*, we can write *∂*_(*l*)_*ϕ_i_*(0,…,0;0) = b and 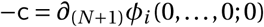. Then eq. (3b) becomes

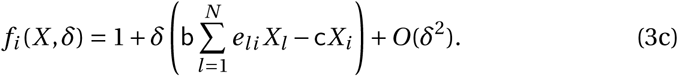

In other words, no matter the choice of the fecundity functions *ϕ*, provided that only the phenotypes of the individuals and their interactants matter, at the first order in *δ* we only need two parameters, b and c, to characterize the fecundity functions.

Our results will be valid for any b and c, but throughout the article, we will consider the case where b > 0 and c > 0, so that type-A individuals are “altruists” providing benefits (b) and paying a cost (c), and we will seek to understand the impact of imperfect strategy transmission on the frequency of altruists.

Finally, we note that when *δ* = 0, all individuals in the population, whichever their type, have the same fecundity: the trait is then neutral.

### 2.3 Reproduction and strategy transmission

The expected number of successful offspring established at site *j* at the next time step, descending from the individual who is living at site *i* at the current time step, is denoted by *B_ji_* (*f* (*X*,*δ*)), written *B_ji_* for simplicity. “Successful offspring” of a focal individual means individuals who descend from this focal individual and who are alive and established at the start of the next time step. Because there is exactly one individual per site, 0 ≤ *B_ji_* ≤ 1. We assume that *B_ji_* does not depend on external factors such as temporal fluctuations independent of the state of the population.

Individuals imperfectly transmit their strategy to their offspring. We do not specify the nature of this transmission (it can be genetic, it can be vertical cultural transmission), but we use for simplicity the term “mutation” to characterize transmission failure. Mutation occurs with probability *μ*, 0 < *μ* ≤ 1; when mutation occurs, the offspring are of type *A* with probability *p* and of type *B* otherwise (0 < *p* < 1). For instance, under this mutation scheme, the offspring of an individual of type *A* is also of type *A* with probability 1 − *μ*+*μp* (Taylor et al., 2007b; Nowak et al., 2010; Tarnita & Taylor, 2014). The parameter *p* controls the asymmetry of mutation, and it is also the expected frequency of type-*A* individuals in the absence of selection (*i.e.*, when *δ* = 0). Although the use of the word “mutation” hints at a genetic transmission of the trait, this framework can also describe vertical cultural transmission, so *μ* does not have to be small. The mutation probability, however, cannot be zero; if it were, the all-*A* and all-*B* states would be absorbing: we would end up either with only type-*A* or only type-*B* individuals in the population, and we would not be able to define a stationary distribution of population states—for similar reasons, *p* cannot be 0 nor 1.

We denote by *D_i_* (*f* (*X*, *δ*)) (or *D_i_* for simplicity) the probability that the individual living at site *i* is dead at the beginning of the next time step, given that the population is currently in state *X*. This probability of death at site *i* can be expressed as a function of the probabilities of birth and establishment of offspring at site *i*, summing over the locations *j* of the potential parents:

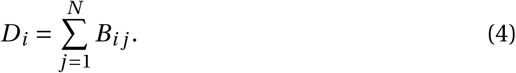

There is exactly one individual per site, so at a given site *i*, there can be at most one successfully established offspring at each time step, and 0 ≤ *D_i_* ≤ 1. On the other hand, the expected number of offspring of the parent currently living at site *i* is 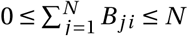.

Finally, we are considering dispersal graphs such that in the absence of selection (*δ* = 0), all individuals have the same probability of reproducing, and all individuals have the same probability of dying—meaning that all sites in the population have the same reproductive value 1/*N* (Taylor, 1990; Caswell, 2001; Lieberman et al., 2005; Maciejewski, 2014; Allen et al., 2015); this implies that for all sites *i*

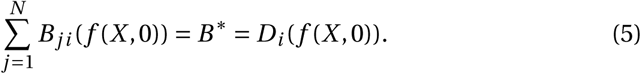

The parameter *B*^*^ is the expected number of offspring produced by an individual during a time step in the absence of selection (*δ* = 0); it is the same for all individuals, but the value taken by *B*^∗^ depends on the life-cycle that is considered.

### 2.4 Life-cycles

Most of our results are derived without specifying a life-cycle (also called “updating rule”). In the *Illustrations* section, we will give specific examples using classical life-cycles: Moran models (Birth-Death and Death-Birth), with exactly one birth and one death during a time step, and the Wright-Fisher model, where all adults die and are replaced by new individuals at the end of a time step.

### 2.5 Simulations

Stochastic simulations, coded in C, were run to numerically confirm the analytical results. For each combination of parameters, a simulation was run for 4 × 10^9^ generations, the state of the population being sampled every 400 generations, where one generation corresponds to *N* time steps with the Moran updating, and 1 time step with the Wright-Fisher updating. A set of parameters corresponded to a choice of updating rule, of population structure, of mutation probability (*μ* ϵ {0.01, 0.025, 0.05, 0.1, 0.15, 0.2}) and of mutation bias (*p* ϵ {0.3, 0.5}.

## 3 Results

### Expected frequency of type-*A* individuals in the population

We describe here the key steps of the computation of the expected frequency of type-*A* individuals in the population and refer the reader to Appendix A for mathematical details.

We denote by Ω the set of all possible states of the population (Ω = {0, 1}^*N*^). No state is absorbing (thanks to mutation, a lost strategy can always reappear), and all states are accessible. We denote by *ξ*(*X*,*δ*,*μ*) the stationary distribution of population states, *i.e.*, the probability that, after a long enough number of time steps, the population is in state *X*, in a model with strength of selection (phenotype differences) *δ* and mutation probability *μ*. Notation 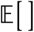 de-notes expectation, for instance the expected state of the population is 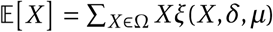. The expected frequency of type-*A* individuals in the population, denoted by 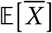, can be computed considering what happens during one during step. Given state *X* of the population, at the end of the time step, the state of the individual living at site *i* depends on whether it has survived during the time step (first term within the brackets of eq. (6)), and, if it has been replaced, on the type of the newly established offspring (second term within the brackets); we then take the expectation over all population states, and obtain:

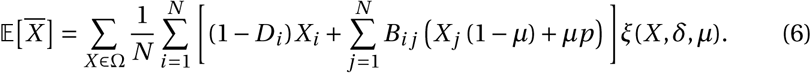

This is the expected frequency of type-*A* individuals in the population. For instance, if we run a simulation of the model for a very long time, the average over time of the frequency of type-*A* individuals will provide an estimation of 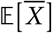; this quantity does not depend on the initial state of the population.

We then assume that selection is weak, *i.e.*, *δ* is small, and write a first-order expansion of eq. (6) that contains derivatives of *ξ*, *D_i_* and *B_ij_* with respect to *δ*. For the last two, we further use the chain rule with the variables *f_k_*, which represent the fecundity of the individual living at site *k*. In doing so, we let appear quantities that are the expectations of the state of pairs of sites when no selection is acting (*i.e.*, when *δ* = 0; we call these “neutral expectations” and *ξ*(*X*,0,*μ*) is called the neutral stationary distribution):

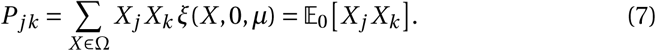

The fact that these neutral expectations appear in our equations does not mean that selection is initially not acting and then “turned on”: selection is acting all the time, but it is weak because phenotypic differences are small (*δ* ≪ 1). At the first order in *δ*, we can ignore the effect of selection on the expected state of pairs of sites, and this is why we only need neutral expectations (eq. (7)).

Eventually, we deduce that the expected frequency of individuals of type *A* in the population can be written as

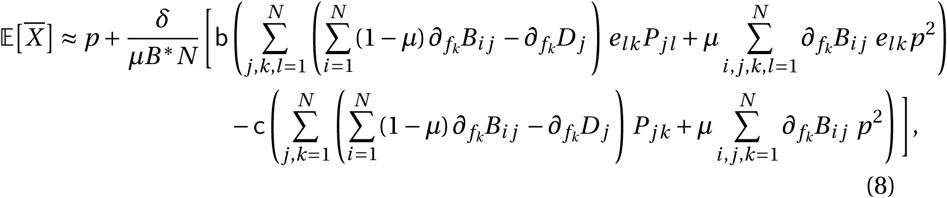

with *P* as defined in eq. (7), *∂_f_k__* being a shorthand notation for 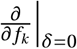, and 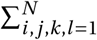 being a compact way of writing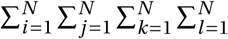. Eq. (8) is an approximation at the first order in *δ* (we neglect terms in *δ*^2^ and higher). A weak mutation approximation of eq. (8) is presented in Appendix A.4.

Eq. (8) is still implicit, because we need to evaluate the *P_ij_* terms, which we now do.

### Expected state of pairs of sites at neutrality

We recall that *P_ij_*, defined in eq. (7), is also the probability that both sites *i* and *j* are occupied by individuals of type *A*, at neutrality (*i.e.*, when *δ* = 0). Under vanishing mutation (*μ* → 0), convenient connections can be made between identity in state and identity-by-descent (Cockerham & Weir, 1993; Rousset & Billiard, 2000), and then with coalescence times (Slatkin, 1991, 1993; Rousset, 2004; Allen et al., 2012). Here as well, we can characterize *P_ij_* in terms of probabilities of identity-by-descent, *Q_ij_*. Two individuals at sites *i* and *j* are said to be identical by descent (IBD) if they share a common ancestor and if no mutation occurred in their lineages since this common ancestor (Kimura & Crow, 1964, note though that the original definition is with an infinite allele model, where each mutation creates a new allele). If two individuals are IBD, then they are both of type *A* with probability *p*, the expected state of a single individual at neutrality. If two individuals are not IBD, then they are both of type *A* with probability *p*^2^. Simplifying, we obtain

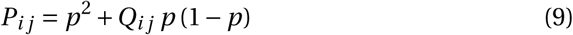

(Rousset & Billiard, 2000; Allen & Nowak, 2014) (see Appendix B.1 for more details). Eq. (9) also valid when *i* = *j*. So we can work with IBD relationships.

To find the probabilities of identity-by-descent, we first write the probability that two individuals at sites *i* and *j* are IBD given the state *X* of the population at the previous time step, and then take the expectation of this conditional probability. We can still do so without specifying the way the population is updated (using notation as in Allen et al. (2015)), and the resulting equation is presented in Appendix B.1, eq. (B.1). This equation can also be adapted to specific updating rules, as shown in the *Illustrations* section (details of the calculations are provided in Appendix B).

Keeping in mind that *Q_ij_* = *Q_ji_* and that *Q_ii_* = 1, we then have to solve a linear system of *N*(*N* – 1)/2 equations to obtain explicit formulas for all the *Q_ij_* terms, for any regular graph. More explicit formulas for *Q_ij_* can be found for regular graphs, and in particular for *n*-dimensional graphs, as we will see in the *Illustrations* section. Finally, we can gather all probabilities of identity by descent in a matrix
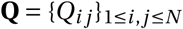.

### Back to the expected frequency of type-*A* individuals

Using the relationship between the expected state of pairs of sites *P_ij_* and probabilities of identity-by-descent *Q_ij_* (eq. (9)), we can rewrite eq. (8) as follows (see Appendix A.5 for details):

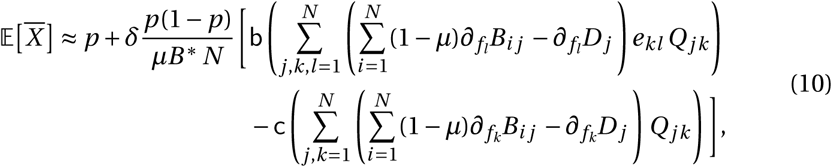

where as before *∂_fk_* is a shorthand notation for 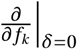, and the sums are written in a compact way.

#### Interpretation

For each focal individual at site *k*, we consider the influence that this individual can have on an identical-by-descent individual at site *j* (*Q_jk_*), by affecting the production of new identical-by-descent individuals by
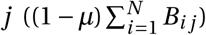, or *j*’s survival (*D*_*j*_). This can occur because of intrinsic changes (the cost of being social c) in the fecundity of individual *k* (*∂_f_k__*), and because the focal *k* provides a benefit to an individual *l* (b *e*_*kl*_) – where *l* is *j* itself or another individual in the population – changing *l*’s fecundity (*∂_f_l__*), with repercussions on *j*. Finally, we note that the factor associated to (−c) is non-negative (see Appendix A.6.)

#### Structure parameter

We say that a strategy is favored if its frequency at the mutation-selection-drift equilibrium is higher than what it would be in the absence of selection. For type *A*, this translates into
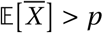. With eq. (10), this condition becomes

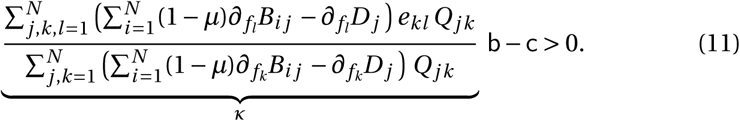

Hence, a single parameter, *κ*, summarizes, for a given life-cycle, the structure of the population and the effect of mutation (Tarnita et al., 2009; Taylor & Maciejewski, 2012); *κ* is interpreted as a scaled coefficient of relatedness, that includes the effect of competition (Lehmann & Rousset, 2010).

#### Alternative formulation

The presence of *μ* at the denominator in eq. (10) might look ominous, given that our equation is meant to be valid for any mutation probability. However, we note that the probabilities of identity by descent can be written

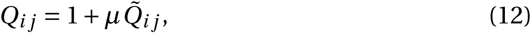

since in the limit *μ* → 0, all individuals in the population are identical by descent. If we now replace *Q_ij_* using eq. (12) in eq. (10), recalling that the size of the population is fixed (eq. (4)), we obtain

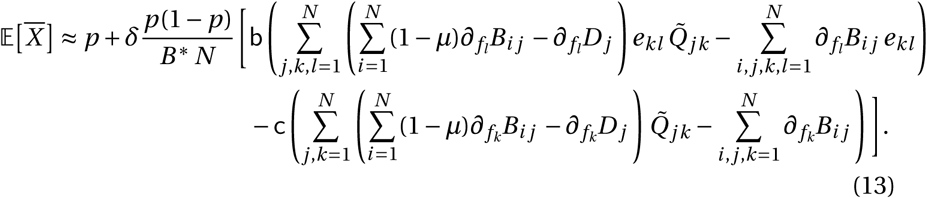

This confirms that dangerous looking denominator *μ* in eq. (10) is not problematic, even for small mutation probabilities. The sums
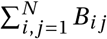
correspond to the total number of births in the population during one time step, which is independent of the composition of the population in the life-cycles that we consider as examples (so the last terms on each line of eq. (13) will disappear).

## 4 Illustrations

### 4.1 Updating rules

The results presented so far were valid for any updating rule, provided it is such that population size remains equal to *N*. We now express the expected frequency of type-*A* individuals for specific updating rules, commonly used in studies on the evolution of altruistic behavior in structured populations: the Moran model and the Wright-Fisher model. Under a Moran model (Moran, 1962), exactly one individual dies and one individual reproduces during one time step; hence, at neutrality, *B*^*^ = 1/*N* (*B*^*^ was defined in eq. (5)). The order of the two events matters, so two updating rules are distinguished (Ohtsuki & Nowak, 2006; Ohtsuki et al., 2006): Birth-Death and Death-Birth. In both cases, payoffs are computed at the start of each time step, before anything happens.

#### 4.1.1 Moran model, Birth-Death

##### Any regular graph

Under a Birth-Death (BD) updating, an individual *j* is chosen to reproduce with a probability equal to its relative fecundity in the population 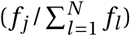; then its offspring disperses at random along the 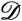 graph, and so replaces another individual *i* with a probability *d_ji_/v*, so that

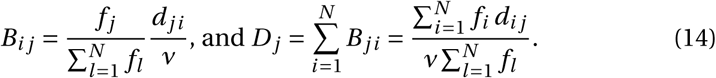

Note that with this updating rule, the probability of dying *D_j_* depends on the composition of the population. With these probabilities of reproducing and dying eq. (10) becomes,

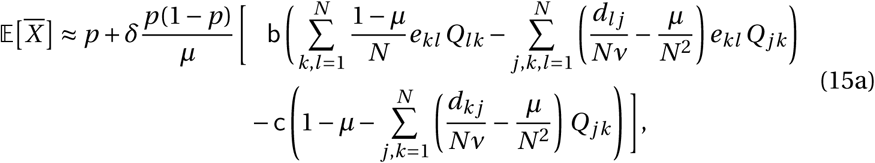

or, using matrix notation,

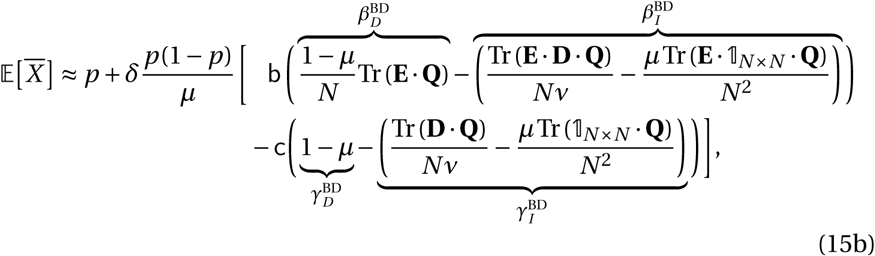

where Tr(**M**) denotes the trace of a matrix **M**, *i.e.*, the sum of its diagonal elements, and 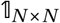 is the *N*-by-*N* matrix of ones. Each of the factors associated to the b and (−c) terms contain direct (_*D*_) effects, discounted by indirect effects (_*I*_). Recall that since we moved from a description with the expected state of pairs of sites (*P_ij_*, eq. (8)) to a description with probabilities of identity-by-descent (*Q_ij_*, eq. (10)), we interpret the different terms in terms of survival and production of identical-by-descent offspring.

##### Interpretation

The direct effect term 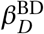 corresponds to the additional identical-by-descent (hereafter, IBD) offspring (1 − *μ*) produced by interacting with IBD individuals 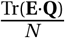 Where there is only one type of interactant (for instance, neighbors on a lattice, or members of the same group) 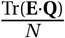 can be described as the relatedness to social interactants times the number of interactants. But when there are different types of interactants (*e.g.*, on a non-symmetric structure like figure 1(a), or when there are weights on the interaction graph ℰ), then we cannot talk of “a” relatedness, and instead consider an averaged relatedness, weighted by the interaction graph ℰ.

Social interactions also have indirect consequences 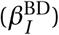. First, a focal *k* that helps (*e_kl_*) an individual *l* who can send offspring (*d_lj_*) to a site *j* occupied by an individual IBD to the focal (*Q_jk_*), indirectly affects the survival of that individual 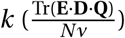; since this is about survival, there is no *μ* involved here. The second term of *β*_*I*_^BD^ corresponds to competitors *l* whose increased fecundity (thanks to interactions with a focal *k*, *e_kl_*) could indirectly reduce the birth rate of individuals *j* IBD to the focal (*Q_jk_*), but whose fecundity increase was “wasted” by the production of non-IBD offspring (*μ*).

The terms associated to (−c) have a similar interpretation. Here, we consider the consequences of the cost of being social, *i.e.*, of the reduction in a focal individual *k*’s fecundity. The direct effect *γ*_*D*_^BD^ corresponds to this reduction of fecundity and its impact on the production of IBD individuals (1 − *μ*). The indirect effects *γ_I_^BD^* are due *i)* to the indirect changes in the survival of an individual *j* IBD to a focal individual *k* (*Q_jk_*), who is less likely to be replaced by the offspring of *k* (*d_kj_*) since *k* is less fecund, and *ii)* to the increased relative fecundity of individuals *j* IBD to the focal *k* (*Q_jk_*), “wasted” by the production of non-IBD offspring (*μ*).

When transmission is almost perfect (*μ* → 0), we recover Grafen & Archetti (2008)’s result that the competition neighborhood under a Birth-Death updating is one dispersal step away (hence the **D** terms in eq. (15b)). Decreasing the fidelity of parent-offspring transmission, by increasing *μ*, not only changes probabilities of identity-by-descent (**Q**), but also the kind of competition to take into account. This is because social interactions affect both the birth and death of individuals, and the issue of transmission fidelity only concerns reproduction, not survival.

##### Probabilities of identity by descent

With the Birth-Death updating rule, the probabilities of identity by descent satisfy, for any *i* and *j* ≠ *i*,

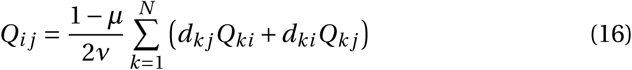

(see Appendix B.2 for details on the derivation). For generic regular graphs, we have to solve a system of *N* (*N* − 1)/2 equations to find the probabilities of identity by descent.

##### Transitive undirected graphs

When the graph is transitive and undirected, probabilities of identity by descent verify

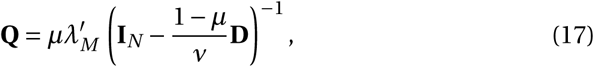

where **I**_*N*_ is the identity matrix, and 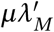 is such that *Q*_*i*_,_*i*_ = 1 for all *i* (the _*M*_ index stands for “Moran”). With eq. (17), eq. (15) simplifies into

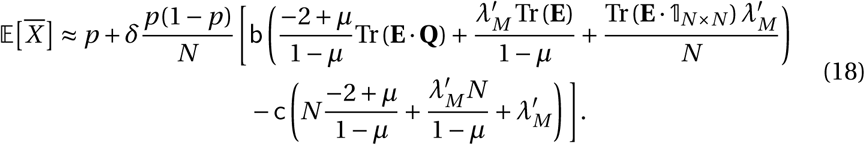

The term Tr(**E**)/*N* corresponds to social interactions with oneself; it is usually considered as null in the case of pairwise interactions, but is not for common good type of interactions (when benefits are pooled and then redistributed). We show in Appendix C.1.3 that the sum of the other two terms associated to the benefits b is negative or zero. So unless interactions with oneself are strong (large Tr(**E**)/*N*), the factor modulating the effect of benefits b is non-positive. We noticed previously that the factor associated to (−c) is non-negative; consequently, the expected frequency of altruists cannot be greater than what it would be in the absence of selection (*i.e.*, 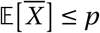.) when interactions with oneself are small.

Evaluating probabilities of identity by descent in transitive regular graphs still requires the inversion of a *N* by *N* matrix (eq. (17)), which can limit applications. Results are simpler in graphs that match our definition of “*n*-dimensional graphs”; they depend on the dimensionality *n* of the graph and are presented in Appendix B.2.

#### 4.1.2 Moral model, Death-Birth

##### Any regular graph

Under a Death-Birth (DB) updating, the individual who is going to die is chosen first, uniformly at random (*i* is chosen with probability 1/*N*). Then, all individuals produce offspring, and one of them (one offspring of parent *j* wins with probability 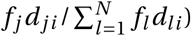 replaces the individual chosen to die. When *d_ii_* ≠ 0, one needs to clarify whether the individual chosen to die reproduces before dying or not; here we assume that this is the case, but some alternative formulations do not. Under this updating rule, we have

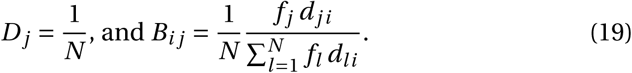

Using matrix notation, eq. (10) becomes

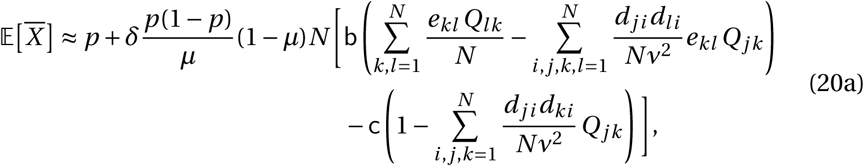

or, using matrix form,

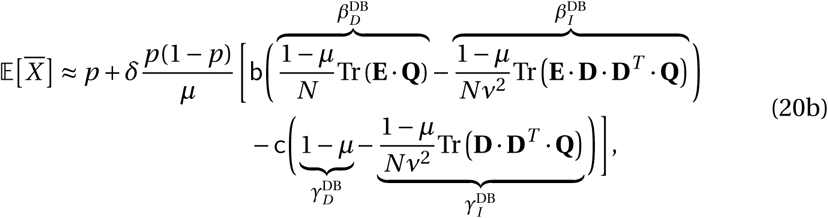

where ^*T*^ denotes transposition.

##### Interpretation

We can again identify direct and indirect effects of benefits and costs; the direct effects (_*D*_) are the same as for the Birth-Death updating rule, but the indirect effects (_*I*_) differ. First, the indirect effects reflect the fact that competitors are now two dispersal steps away (Grafen & Archetti, 2008; Débarre et al., 2014). Under a Death-Birth updating rule indeed, individuals *j* and *k* are competing for a site *i* whose occupant has just been chosen to die if both *j* and k can send their offspring to *i*; this depends on *d_ji_d_ki_*, leading to the **D** · **D**^*T*^ products in eq. (20). Second, with a Death-Birth updating, social interactions do not affect the probability of dying, so we only take into account effects on reproduction, and we can factor in the (1 − *μ*) terms.

##### Probabilities of identity by descent

With the Death-Birth model as defined above, the system of equations for the probabilities of identity by descent at neutrality is the same as in eq. (16).

##### Transitive undirected graphs

When the graph is transitive and undirected, eq. (17) still holds and eq. (20) simplifies into

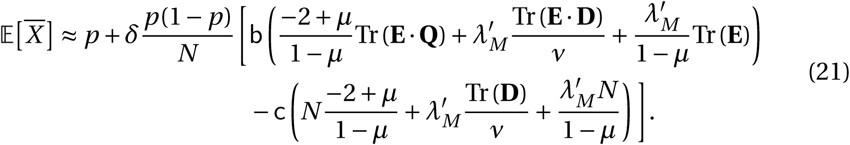

Now, even in the absence of self-interactions (*i.e.*, even when Tr(**E**) = 0), the term associated to b can be positive. As it is the case in the absence of mutation (Ohtsuki et al., 2007; Taylor et al., 2007a; Débarre et al., 2014), a key role is played by 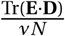, which is higher the more the *D* and *ℰ* graphs overlap; if we also scale the interaction graph *ℰ* to control for the number of interactants, then a lower degree makes 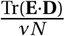 higher, and thereby increases the expected frequency of altruists in the population.

We also note that eq. (21) (Death-Birth) and eq. (18) (Birth-Death) become the same when 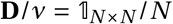, *i.e.*, when the dispersal graph is the complete graph (with self-loops), in other words when the population is unstructured. This reflects the fact that in the Birth-Death updating, the individual who reproduces is chosen among all individuals of the population, while in the Death-Birth updating, the individual who reproduces is chosen locally, among the neighbors of the individual who just died. This scaling persists with arbitrary fidelity of parent-offspring transmission *μ*.

#### 4.1.3 Wright-Fisher

Under a Wright-Fisher model, generations are non-overlapping: all adults produce offspring, then all adults die and the offspring disperse and compete for establishment, so that

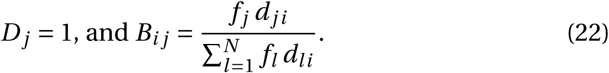

In a Wright-Fisher model, at neutrality, *B*^*^ = 1 (the entire population is renewed at each generation; in a Moran model we had *B*^*^ = 1/*N*); eq. (22) differing from its Moran Death-Birth equivalent (eq. (19)) by only a factor 1/*N*, we end up with the same equation as eq. (20) for the expected frequency of type-*A* individuals in the population. The difference between the Moran Death-Birth and Wright-Fisher life-cycles however lies in the evaluation of probabilities of identity by descent.

##### Probabilities of identity by descent

Under a Wright-Fisher model, the entire population is replaced, so the equation is different from the one obtained under a Moran model; probabilities of identity by descent of two different individuals satisfy (*i* ≠ *j*)

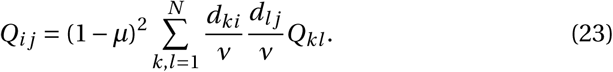

(see Appendix B.3 for details of the derivation.) In short, two individuals are identical by descent if there parents were, and if neither offspring is a mutant ((1 − *μ*)^2^).

##### Undirected transitive graphs

When the dispersal graph is undirected (**D** = **D**^*T*^) and transitive, the probabilities of identity by descent verify

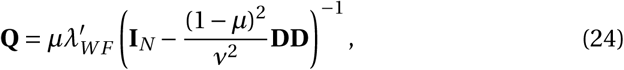

with 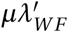 such that for all *i*, *Q_ii_* = 1, and the _*WF*_ index stands for “Wright-Fisher”. With this, the expected frequency of type-*A* individuals becomes

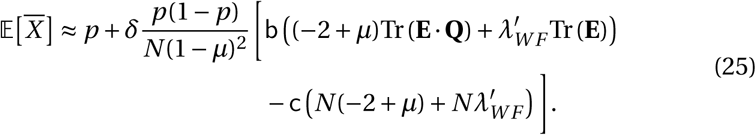

We can immediately see the difference with the Moran Death-Birth case (eq. (21)), caused by a different equation for the probabilities of identity by descent **Q**. Crucially missing in eq. (25) is the positive term 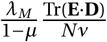: without it, the factor associated to the benefits b is negative unless interactions with oneself (Tr(**E**)) are strong enough, as was the case with the Moran Birth-Death updating.

As for the Moran model, evaluating probabilities of identity by descent in undirected transitive graphs (eq. (24)) involves the computation of the inverse of a *N* by *N* matrix. More explicit results can be obtained for “*n*-dimensional graphs”; they are presented in Appendix B.3.

### 4.2 Specific population structures

All numerical examples given in this section are derived with b > 0 and c > 0, so type-*A* individuals can be called altruists.

As an illustration, we explore the impact of mutation on the expected proportion of type-*A* individuals in graph-structured populations, in which the same graph defines dispersal and interactions among individuals (Lieberman et al., 2005; Hindersin & Traulsen, 2015; McAvoy & Hauert, 2015), so that **E** = **D**.

When the graph undirected and transitive, the equations for the expected frequency of altruists (type-*A* individuals) can be further simplified; the formulas are given in the Appendix(eq. (C.8) and eq. (C.12)). Under a Wright-Fisher updating, eq. (25) cannot be much further simplified.

### 4.2.1 Small graphs

For regular graphs of small size, the probabilities of identity by descent can be calculated directly using eq. (16) (Moran model) or eq. (23) (Wright-Fisher). In figure 2, we show the value of 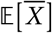 on three regular graphs that have the same size (*N* = 12) and the same degree (*v* = 3), and we consider three common life-cycles in populations of fixed size (Moran Death-Birth, Moran Birth-Death, Wright-Fisher). We compare the prediction based on eq. (8) (curves) to the outputs of stochastic simulations (points) (Comparable results are obtained with other values of mutation bias *p*, see figure S1). For all life-cycles, increasing the mutation probability *μ* makes 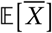 closer to its value at the mutation-drift equilibrium (*p*). The curves corresponding to different structures are almost undistinguishable under a Moran model (figures 2(a) and (b))—the curve corresponding to the graph with no symmetry (red, squares) being a bit less similar though). In the Wright-Fisher model (figure 2(c)) however, the effects of the three structures are clearly different, even when *μ* becomes very small: knowing only the size (*N*) and degree (*ν*) of a regular graph is not enough in this case to precisely predict the expected frequency of altruists in the population. This is because the *λ^’^_WF_* terms greatly differ between the three graphs that we tested, all the more when *μ* → 0, while the values of 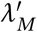 for the three structures remained close to each other.

### 4.2.2 Large graphs: variations on a circle

When the number of nodes gets larger, we have to concentrate on graphs with a high level of symmetry. Here we will consider 1-dimensional graphs (graphs whose nodes can be relabelled to satisfy eq. (2)) that are undirected, and hence that can be categorised as undirected transitive graphs. For simplicity, we can consider a circle graph, such that the nodes are arranged on a circle, and each node is connected to its two neighbors only. Here, we assume that the number of nodes is infinite: *N* → ∞. As previously, a given node hosts exactly one individual (see figure 3(a)).

Under a Moran model, using eq. (B.12b), we find for *μ* > 0

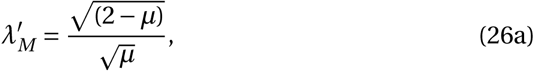

and, although the quantity is not needed to compute 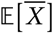 under a Moran model (see eq. (C.8) and eq. (C.12)), the probability of identity by descent between two neighbors on the circle is given by

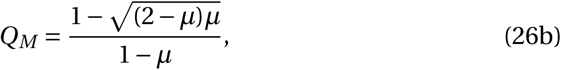

and we recover the formula presented in, *e.g.*, Allen et al. (2012) (see Appendix B.2.4 for details). This result is plotted in figure 3(c). We however need to note that the first-order approximation for 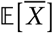 fails when both *μ* → 0 and *N* → ∞: this is because the integral behind eq. (26a) does not converge when *μ*→ 0. Similarly, for instance, the first order approximation for the probability that two neighbors are identical by descent 1 − *μ*(*N* − 1), which was obtained by Taylor et al. (2007a), fails when *N* is too large compared to *μ*.

Under a Wright-Fisher updating, the probability of identity by descent between neighbors is equal to 0. This is because all individuals reproduce at each time step, and their offspring can only establish on the node on the left or on the right of their parent, so that relatedness cannot build up (a feature called checkerboard effect by Grafen & Archetti, 2008). This checkerboard effect is also the reason why 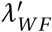 differed among the small graphs that we tested; for instance, under a Wright-Fisher updating **Q** does not converge to 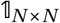 when *μ* → 0 with the graph depicted in figure 1(c) while it does for graph 1(b).

We can however modify the graph to allow for establishment in the parent’s node: with probability (1 − *m*) the offspring remain where the parent was, otherwise they move to the right or the left-hand side node (with probability *m*/2 for each; see figure 32(b)). In this case, we find the following probability of identity by descent between neighbors:

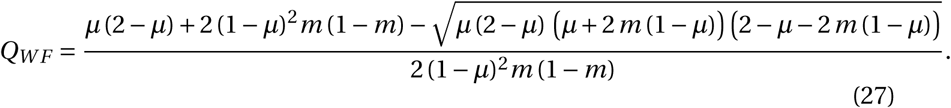

(See Appendix B.3.4 for details; the corresponding value of 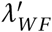 is given in eq. (B.48b).) *Q*_*W F*_ is undefined for *μ* = 0 or *m* = 1, and lim_*μ*→0_ *Q_WF_* = 1, but lim_*m*→0_ *Q_WF_* = 0. The result is plotted in figure 3(d) for different values of the emigration probability *m*.

## 5 Discussion

While most studies on the evolution of cooperation assume an almost perfect fidelity of strategy transmission from parent to offspring, here, we explored the effect of arbitrary mutation on the evolution of social behavior in structured populations. We provide a formula (eq. (10)) that gives the expected frequency of a given strategy, for any life-cycle, any fidelity of parent-offspring strategy transmission, and that is valid in populations of fixed size that are such that the reproductive values of all sites are equal (*i.e.*, when all individuals have the same fecundity, they all have the same chance of actually reproducing). The formula depends on the probability of identity by descent of pairs of individuals, and we show how to compute those in general.

### Identity by descent and expected state of pairs of sites

The effects of social interactions depend on the actual types of the individuals who interact. With imperfect strategy transmission from parents to their offspring (*μ* > 0), common ancestry does not guarantee that two individuals are of the same type. The concept of identity by descent, as we use it in this article, adds to common ancestry the condition that no mutation has occured in the two individuals’ lineages since the common ancestor (Kimura & Crow, 1964; Taylor et al., 2007b), and hence guarantees that the two individuals are of the same type. Two individuals that are not IBD can be treated independently, and we can hence relate the probability that the individuals at two sites *i* and *j* to their expected state (see our eq. (9), Allen & Nowak (2014), or also Rousset & Billiard (2000)). Finally, equations with probabilities of identity by descent are simpler than those for the expected state of pairs of sites.

### A structure parameter *κ*

Tarnita et al. (2009) and Taylor & Maciejewski (2012) showed that, when social interactions affect fecundities, there exists a parameter independent of the terms of the interaction matrix that summarizes the effects of population structure (in terms of dispersal patterns and also of who interacts with whom), that depends on the rule chosen to update the population and on mutation; here we provide a generic formula for such a structure parameter. This parameter, *κ*, can be interpreted as a scaled relatedness (Queller, 1994; Lehmann & Rousset, 2010), which includes the effect of competition. Eq. (11) provides a generic formula for *κ*, for any life-cycle and population structure (provided condition eq. (1) is satisfied).

The actual value of the scaled relatedness *κ* depends on the life-cycle and on the mutation probability *μ*. First, *κ* includes competition (what we call “indirect effects” of social interactions), and the scale of competition depends on the life-cycle (Grafen & Archetti, 2008; Débarre et al., 2014). Second, even direct effects of social interactions—and so even what is referred to as relatedness—do depend on the life-cycle and *μ*.

Finally, there is a single structure parameter *κ* because social interactions only affect fecundity. Previous studies assuming vanishing or absent mutation have shown that the parameter will be different if social interactions instead influence survival (Nakamaru & Iwasa, 2006; Taylor, 2010) and that we need more than one parameter if social interactions affect both fecundity and survival (Débarre et al., 2014).

### Updating rules and the evolution of altruism

We illustrate our results with specific updating rules, with either exactly one new individual at each time step (Moran Birth-Death, Moran Death-Birth), or exactly *N* new individuals, *i.e.*, the entire population being renewed at each time step (Wright-Fisher). Previous studies done under the assumption of vanishing mutation rates (and with undirected transitive dispersal graphs) found that updating rules had a great impact on the evolution of altruism, and in particular, that selection did not favor altruism (benefits given to others exclusively) under a Wright-Fisher or Moran Birth-Death updating (the “cancellation result”; Taylor, 1992; Taylor et al., 2011; Ohtsuki et al., 2007; Lehmann et al., 2007). The result holds with imperfect strategy transmission as well. Under a Death-Birth updating, competition is against individuals two dispersal steps away, but identity by descent is computed using individuals one dispersal step away: competition is “diluted”, and altruism can be favored by selection.

To include imperfect transmission in our model, we had to distinguish between effects of social interactions on the probability of reproduction (*∂_fl_B_ij_*; because mutation can occur) or of death (*∂_fl_D_j_*; transmission fidelity is not relevant). Thanks to this, we can trace the origin of the indirect effects, and we can highlight the different nature of the competitive circles identified by Grafen & Archetti (2008), for the Birth-Death and Death-Birth models, in top of their differences in radii. Under a Birth-Death model, with vanishing mutation, the competitive circle is a circle of death: if a focal individual has an increased fecundity, this reduces the survival of its neighbors. Under a Death-Birth model however, the competitive circle is a circle of (reduced) birth: if a focal individual has an increased fecundity, this reduces the reproductive potential of its neighbors’ neighbors.

Again, note that the conclusions for the Moran model depend on which trait is affected by the social behavior. If survival, instead of fecundity, is affected by the social behavior, the scale of competition (Grafen & Archetti, 2008) is switched between the Birth-Death and Death-Birth updating rules, and then altruism is favored under a Birth-Death updating (Nakamaru & Iwasa, 2006; Taylor, 2010; Débarre et al., 2014).

### Implications for adaptive dynamics

Our results are obtained by considering the changes that occur during one time step from a given population state, chosen from the stationary distribution of population states—hence the phrase “long-term”, which differs from the use made by, for instance Van Cleve (2015), where it refers to a trait substitution sequence. Yet, our results can also be used in that context. The adaptive dynamics framework describes evolution as a series of trait substitutions (Geritz et al., 1997; Champagnat et al., 2006; Champagnat & Lambert, 2007; Lehmann, 2012; Lehmann & Rousset, 2014) and is based on the assumption that mutations are rare and incremental; in a finite population, trait evolution proceeds along a gradient of fixation probabilities. Computing those fixations probabilities can be challenging in spatially structured populations.

Yet, the existence of a single parameter (in this case, defined as *σ* = (*κ* − 1)/(*κ* + 1), Tarnita et al., 2009) to characterize population structure and update rules led to the extension of the adaptive dynamics framework to populations with arbitrary structure (Allen et al., 2013), the structure parameter however remaining unspecified in general. Our formula for *κ* (eq. (11)) is valid for arbitrary mutation, so *a fortiori* for vanishing mutation probabilities, and can therefore be used to explicitly study adaptive dynamics in structured populations (provided the reproductive values of all sites are equal).

## Data accessibility

All codes are provided as a zipped folder, and can be downloaded from https://figshare.com/articles/Mutation_and_social_evolution/3207748. They will be uploaded on Dryad when the manuscript is accepted.

## Acknowledgements

I thank Sally Otto and François Bienvenu for comments on the manuscript and Ben Allen for clarifications. The comments and suggestions of Alan Grafen and an anonymous reviewer helped improve the manuscript. The simulations were run on the Migale cluster (http://migale.jouy.inra.fr/).

## Appendix A

### A Expected frequency of type-*A* individuals

#### A.1 Conditional expectations

We denote by 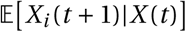the expected state of the individual at site *i* at time *t* + 1, given that the population is in state *X* at time *t*. Because *X*_*i*_ is an indicator variable, 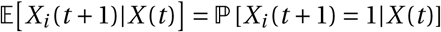. Site *i* is occupied by an individual of type *A* at time *t* + 1 if: *i)* it was occupied by an individual of type *A* at time *t* and this individual has not been replaced (*i.e.*, has not died) between *t* and *t* + 1 (first term in eq. (A.1)), or *ii)* the individual has been replaced by a new one, whose parent was in site *j* at *t*; in this case, either the parent was of type *A* and the offspring is not a mutant; or, whichever the type of the parent, the offspring is a mutant and mutated into type *A* (second term of eq. (A.1)):

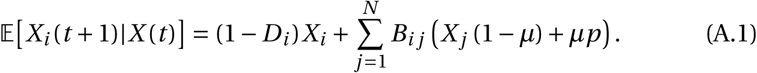

#### A.2 Unconditional expectations

We now want to consider the long-term outcome of competition. We denote by *ξ*(*X,δ,μ*) the probability that the population is in state *X,* given phenotype difference *δ* between the two types and a mutation rate *μ*, and by Ω the ensemble of all possible population states. By definition, the expectation of the state of the population is given by 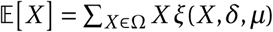.

When the station distribution is reached (*i.e.*, for very large *t*), 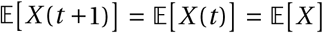; we consider the population average of *X*, 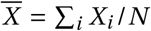. From eq. (A.1), we obtain

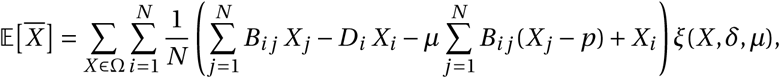

which, after simplifications, becomes

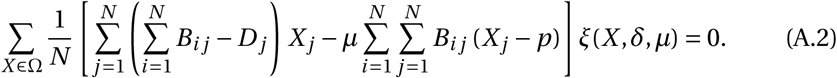

##### Weak selection approximation

While eq. (A.2) is valid for any *μ* and *δ*, we now assume that *δ*, which scales the phenotype difference, is small, so that we can neglects terms of order *δ*^2^ and higher. We note that in the absence of selection (*i.e.*, when the expressed phenotypes are identical, *δ* = 0), the expected state of a site *j* when the stationary distribution is reached is equal to the probability that a mutated offspring is of type *A* (*i.e.*, 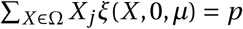; see section A.3 below for more details). Using eq. (5) and the compact notation *∂_δ_* to represent 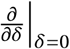, a first-order expansion of eq. (A.2) yields, after simplifications:

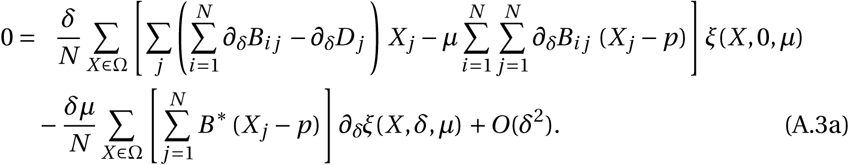

Because *ξ* is a probability distribution, 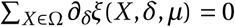 reorganizing eq. (A.3a), we obtain

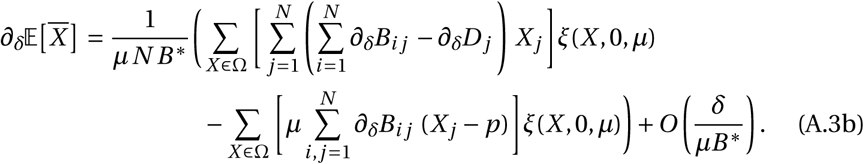

We can now use the chain rule:

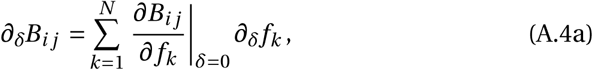

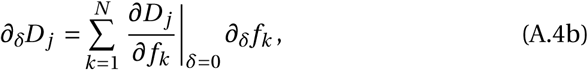

where the *∂_δ_f_k_* terms are computed using the definition of *f* presented in eq. (3c):

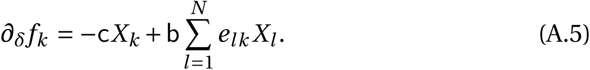

Doing so, we let products of *X* s appear. We denote by *P_jk_* the expected state of a pair of sites (*j*, *k*) evaluated when there are no social interactions (*δ* = 0):

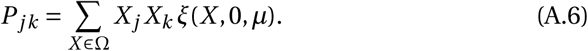

Combining these results and plugging them in the first order expansion of 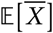,

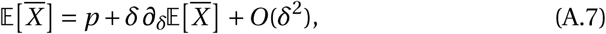

we recover eq. (8) in the main text.

#### A.3 In the absence of selection (*δ* = 0)

In the absence of selection, neither *D*_*i*_ nor *B*_*ij*_ depend on the state of the population, because all individuals now have the same fecundity; we then denote them by *D*_*i*_^0^ and *B*_*i*__*j*_^0^. Consequently, when *δ* = 0, and given that neutral reproductive values are all equal (eq. (5) in the main text), eq. (A.1) becomes

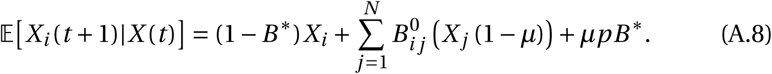

We now take the expectation of eq. (A.8) over the neutral distribution of states (*ξ*(*X*,0,*μ*); we denote by 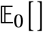 this neutral expectation); since 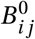 does not depend on *X*, we have

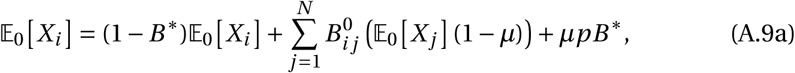

and we obtain after simplifying

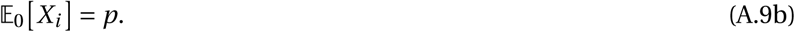

#### A.4 Weak mutation

When *μ* = 0, there is no stationary distribution of states, because the states *X* = **0** and *X* = **1** (loss of type-*A* and loss of type-*B* individuals, respectively) are absorbing. We can nevertheless extend *ξ* by continuity at *μ* = 0, so that *ξ*(*X*,*δ*,0) = lim_*μ*→0_ *ξ*(*X*,*δ*,*μ*). Then, it does not matter whether we Taylor-expand *ξ* first in *δ* then in *μ* or first in *μ* and then in *δ*, and so we can consider *μ* ≪ *δ* and *δ* ≪ *μ* (Tarnita & Taylor, 2014).

##### Weak selection then weak mutation

Starting from eq. (A.3a), a first order expansion near *μ* = 0 yields

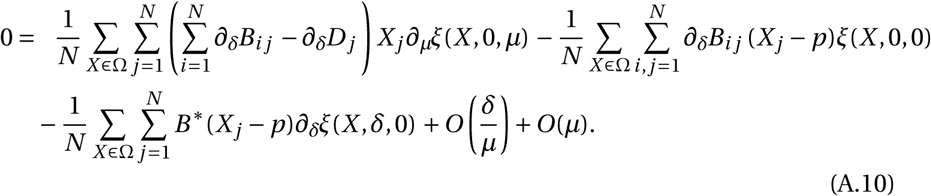

Here we have *δ* ≪ *μ* ≪ 1. Notation *∂*_*μ*_ stands for 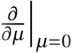.

##### Weak mutation then weak selection

Starting from eq. (A.2), a first order expansion near *μ* = 0 and then a first order expansion near *δ* = 0 yields

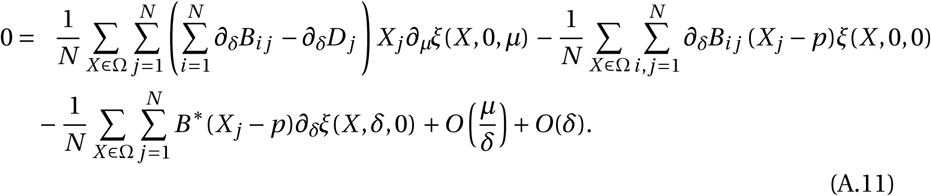

Here we have *μ* ≪ *δ* ≪ 1.

At the first orders, eq. (A.10) and eq. (A.11) are the same.

When *μ* → 0, the population is either in state *X* = **0** or in state *X* = **1**, so

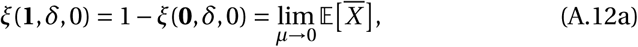

and as a result

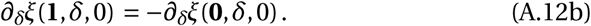

In addition, when *δ* = 0,

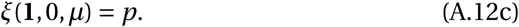

So at the first orders, reorganizing eq. (A.10) (or equivalently eq. (A.11)), we obtain the following equation for the derivative with respect to *δ* of the expected state of the population when *μ* → 0 (Tarnita & Taylor, 2014):

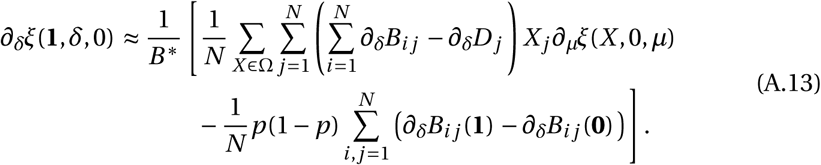

#### A.5 With probabilities of identity by descent

Here we detail how to go from eq. (8) to eq. (10). We use the relationship with *P*_*ij*_ and *Q*_*ij*_ given in eq. (9) (see Appendix B for more details on how to obtain this relationship), and obtain

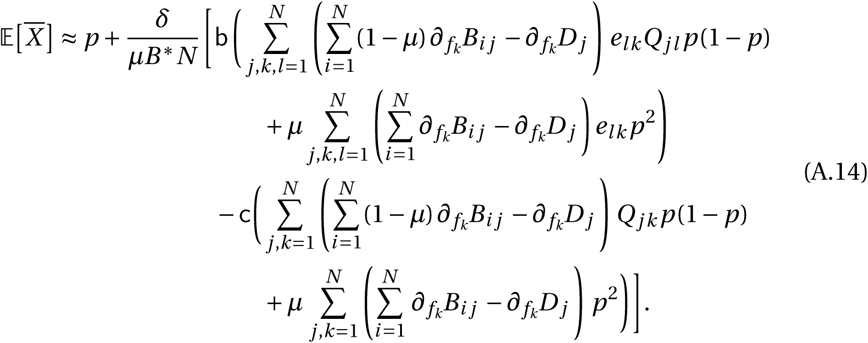

Since the size of the population is constant (eq. (4)) 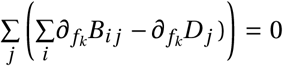 and the second terms of each factor in eq. (A.14) are zero, and we obtain eq. (10) in the main text (where we switch the names of the summation indices *k* and *l* for the factor associated to b).

#### A.6 Sign of the factor associated to (−c)

Eq. (10) is holds for any combination of parameters, and in particular, the factors associated to b and (−c) do not depend on b and c, but depend on population structure (via the dispersal and interaction graphs), mutation probability *μ* and on the chosen updating rule. So in particular when b = 0 and c > 0, eq. (10) reduces to

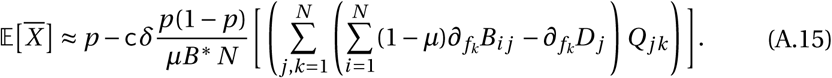

Having b = 0 and c > 0 means that there are no social interactions between individuals, and that type-*A* always have a lower fecundity than type-*B* individuals. For any type of structure, mutation probability and updating rule, the expected frequency of type-*A* individuals is hence lower that what it would be when *δ* = 0, which means that the factor associated to (−c) is non-negative.

## Appendix B

### B Probabilities of identity by descent

We first start by showing the link between the expected state of a pair of sites (*P*_*ij*_) and probabilities of identity by descent (*Q*_*ij*_), for any life-cycle.

#### B.1 Any life-cycle

##### B.1.1 Notation

To be able to consider any life-cycle, we use notation similar to what is used in Allen et al. (2015). At each time step, from 1 to *N* individuals are replaced, depending on the updating rule; *R* denotes the set of individuals that are replaced (*i.e.*, the sites where an individual is replaced by another one). For each site *i* where a replacement happened (*i* ∈ *R*), *α*(*i*) gives the index of the site where the parent of the new individual lived, while for individuals that were not replaced, ∀*i*∈ {1, …, *N*}\*R*, *α*(*i*) = *i*^1^. Finally, *ρ*(*R*, *α*) denotes the probability of the replacement event (*R*, *α*). In the absence of selection, this probability does not depend on the current state of the population.

##### B.1.2 Expected state of a pair of sites

Considering two different sites *i* and *j*, depending on the updating rule, at each time step, *i*) either none of the individuals are replaced—then they are both of type *A* if they already were [first term in eq. (B.1)], *ii*) either one of the individuals (*i* or *j*) is replaced—then they are both of type *A* if the surviving individual is *A* and if either the parent of the other individual was of type *A* and no mutation occurred, or the offspring mutated into type *A* whichever the type of its parent [second and third terms in eq. (B.1)]), or finally *iii*) both individuals are replaced—then the probability that both offspring are of type *A* is 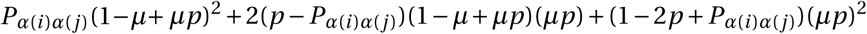, which simplifies into the fourth term in eq. (B.1)). We obtain the following equation:

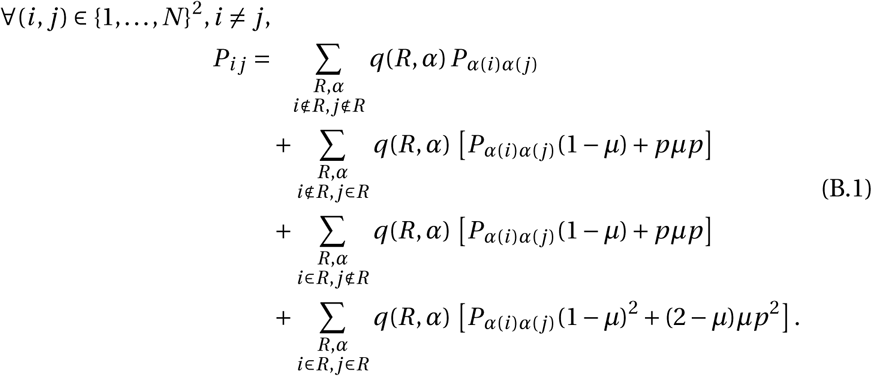

##### B.1.3 Identity by descent

Considering two different sites *i* and *j*, depending on the updating rule, at each time step, *i*) either none of the individuals are replaced—then they are identical by descent (IBD) if they already were [first term in eq. (B.1)], *ii*) either one of the individuals (*i* or *j*) is replaced—then they are both IBD if the surviving individual and the parent of the new individual were and no mutation occurred [second and third terms in eq. (B.1)]), or finally *iii*) both individuals are replaced—then then are IBD if their two parents were and no mutation occurred in either [fourth term in eq. (B.1)]. We obtain the following equation:

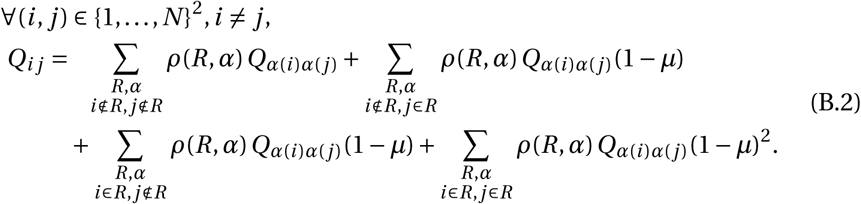

For all pairs *i* ≠ *j*, eq. (B.1) and eq. (B.2) are equivalent when we set

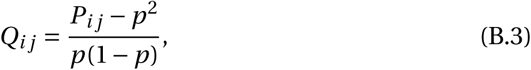

and eq. (B.3) is also valid when *i* = *j* (in this case *Q*_*ii*_ = 1 and *P*_*ii*_ = *p*). So we can use the recursion on *Q* presented in eq. (B.2) together with eq. (B.3).

Finally, while *Q*_*ij*_ is an expectation over the stationary distribution of population states, we also introduce the indicator variable *q*_*ij*_ (*t*), equal to 1 if, in a realization of the process, the individuals at sites *i* and *j* are IBD at time *t*. We also denote by **Q** the matrix gathering the *Q*_*ij*_ terms.

#### B.2 Moran model

In a Moran model, exactly one individual died and one individual reproduces during one time step. Given a state *X* at time *t*, for *i* ≠ *j*, probabilities of identity by descent verify

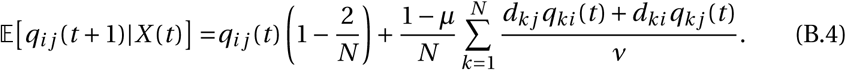

Taking the expectation of this quantity over the stationary distribution of states, we obtain

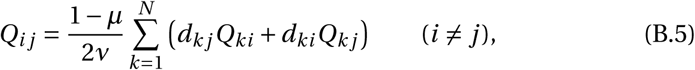

and *Q*_*ij*_ = 1 when *i* = *j*. Eq. (B.5) is valid for any regular graph; all the *Q*_*ij*_ terms can be found by solving a system of *N* (*N* − 1)/2 equations (since *Q*_*ij*_ = *Q*_*ji*_). We can also write eq. (B.5) in matrix form:

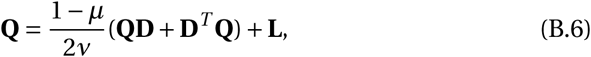

where **D** is the adjacency matrix of the dispersal graph (with elements *d*_*ij*_), ^*T*^ denotes transposition, and **L** is a diagonal matrix whose *i* th diagonal element is 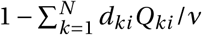
*i.e.*, such that *Q_ii_* = 1). We note that each of these elements is of order *μ* (because when *μ* → 0, 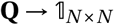, and then 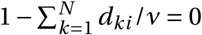 since the graph is regular).

##### B.2.1 Transitive undirected graphs

When the dispersal graph is transitive, then all the elements on the diagonal of **L** are equal, so we can write 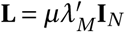 where **I**_*N*_ is the *N* by *N* identity matrix. When the graph is also undirected, **D** = **D**^*T*^, and we also show by induction that **DQ** = **QD** (Grafen & Archetti, 2008).

Let us assume without loss of generality that initially (*t* = 0) all individuals are IBD (*q*_*ij*_ (0) = **1**_*N N*_, where **1**_*N N*_ is the *N*-by-*N* matrix containing only ones) and of type *B* (*X* (0) = {0,…,0}). Also, let us denote by *ζ*_0_(*X*, *t*) the probability that the population is in state *X* at time *t* given that it was in state {0,…,0} at time 0, and by 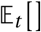 expectations with respect to that distribution, at time *t*. Then from eq. (B.4), since *q*_*ii*_ = 1, and given that the graph is regular,

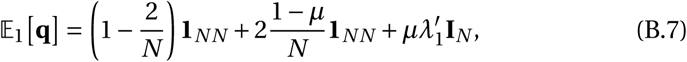

so

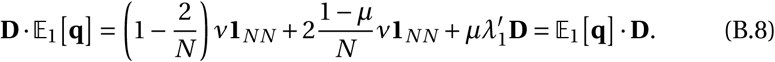

Then, assuming that **D** and 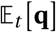 commute, and given that we assume **D** = **D**^*T*^,

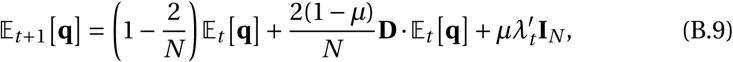

so

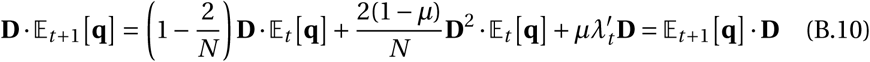

And so, when *t* → ∞, we have **D** · **Q** = **Q** · **D**.

Then with a transitive undirected dispersal graph, eq. (B.6), simplifies into

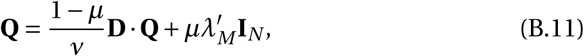

and so (for *μ* > 0),

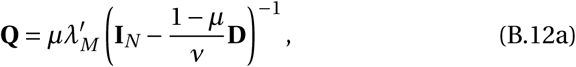

with

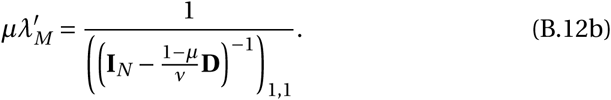

Eq. (B.11) also implies

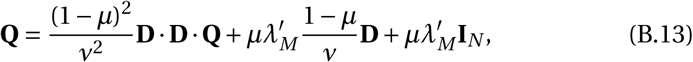

but also

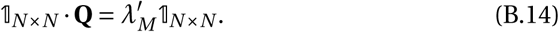

###### More about 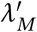

We denote by **u** the column vector of ones; then from eq. (B.11) we have, using the fact that **D** and **Q** commute,

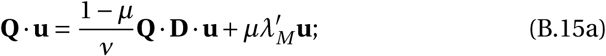

since the dispersal graph is regular, we obtain

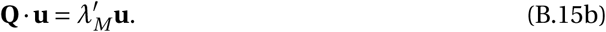

So *λ*′_*M*_ is an eigenvalue of **Q**. When *μ* → 0, **Q** → 𝟙_*N*×*N*_, and eq. (B.15b) implies that

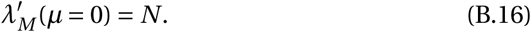

In the main text, we decomposed probabilities of identity by descent as *Q*_*ij*_ = 1 + *μQ*_*ij*_ (eq. (12)); since we are dealing with probabilities, we have 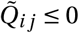 for all *i* and *j* (it is equal to 0 when *i* = *j*). Similarly, using eq. (B.16) we write

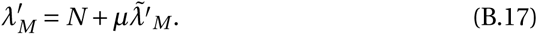

Putting this back into eq. (B.15b), we obtain

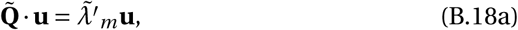

*i.e.*,

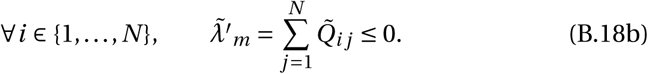

Altogether, this means that

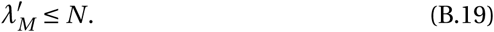

It is possible to find more explicit formulae when the graphs are transitive and when they are *n*-dimensional, and we do so for 1-D and 2-D graphs.

##### B.2.2 One-dimensional graphs

On a 1-D graph, numbering the different nodes modulo *N*, for all *i* and *j*, by definition of a 1-D graph, 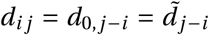, and as a result similar equalities hold for the expected states of pairs of sites: 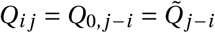. We can hence rewrite eq. (B.5) as follows, keeping in mind that *Q*_*ij*_ = *Q*_*ji*_ and that node numbering is done modulo *N*:

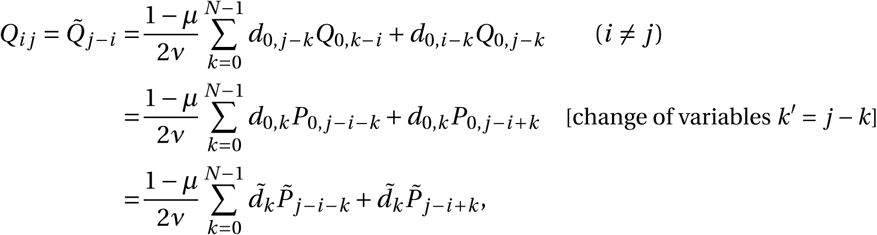

so that

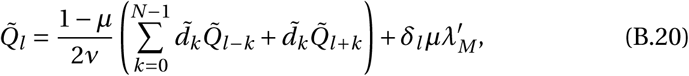

where *δ*_*l*_ = 1 when *l* ≡ 0 and *δ*_*l*_ = 0 otherwise, and *λ* is as defined in the previous section, *i.e.*, such that 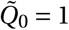:

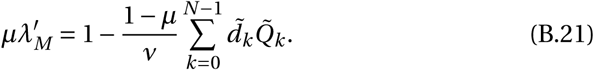

(Recall that 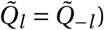.

To solve for 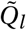, we can follow the same method as in Malécot (1975); Gandon & Rousset (1999) and use discrete Fourier transforms, defining the transforms of *Q* and of *d* as follows:

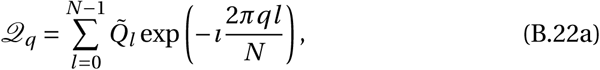

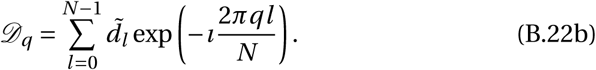

and in particular (*ν* being the degree of the dispersal graph)

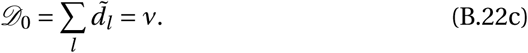

We obtain

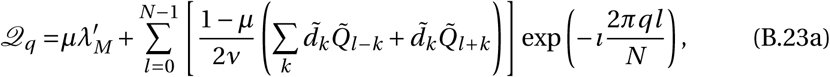

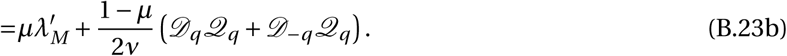

Solving for 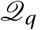, we obtain

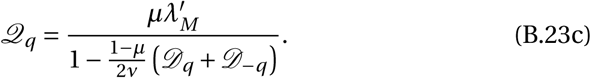

To recover 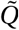, we now use an Inverse Discrete Fourier Transform

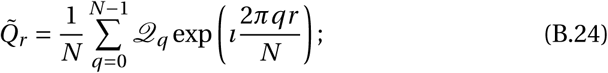

combining eq. (B.23c) and eq. (B.24), we obtain

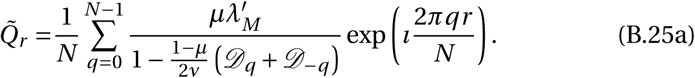

When *r* = 0, we have 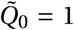, so combining this with eq. (B.25a), we can now evaluate *λ*:

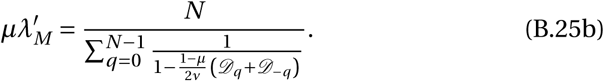

Finally, when the graph is not oriented, 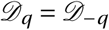.

##### B.2.3 Two-dimensional graphs

Similar calculations are done with two-dimensional graphs. Numbering is done modulo *N*_1_ for the first dimension, and modulo *N*_2_ for the second dimension (*N*_1_*N*_2_ = *N*). The 2-D equivalent of eq. (B.20) is

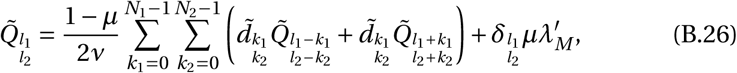

where 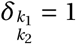 when (*k*_1_, *k*_2_) ≡ (0, 0) (modulo *N*_1_ and *N*_2_), and 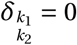 otherwise, and

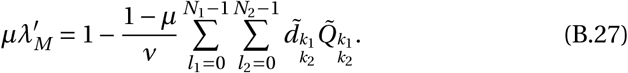

We then use 2-D Discrete Fourier Transforms:

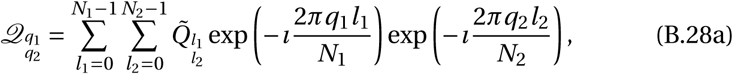

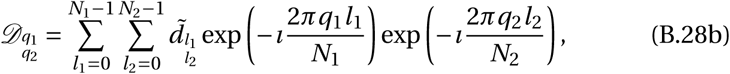

and obtain

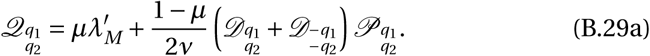

Solving for 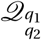,

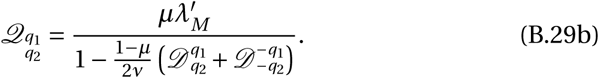

Finally, an Inverse Fourier Transform gives us 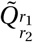:

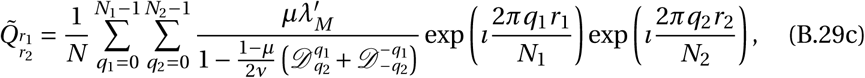

with *C* such that 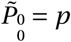:

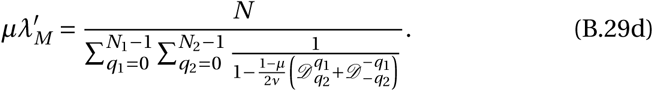

And when the graph is undirected, 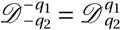.

##### B.2.4 Illustration: infinite circle

On a circle graph (like in figure 3(a)), the Fourier transform of the dispersal distance is

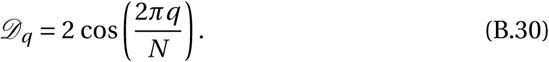

We can evaluate 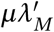 using eq. (B.25b),

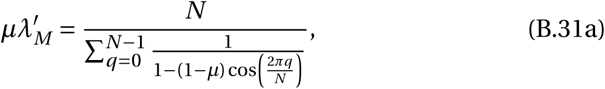

and when population size is infinite, this becomes

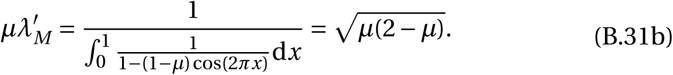

But we note that the integral does not converge when *μ* → 0. Finally, we compute probabilities of identity by descent using eq. (B.25a), and obtain eq. (26b) in the main text for neighbors on the the circle (*q* = 1).

#### B.3 Wright-Fisher model

In a Wright-Fisher model, all individuals are replaced at each time step. Given a state *X* at time *t*, for *i* ≠ *j*, probabilities of identity by descent verify

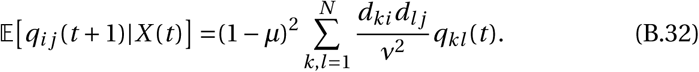

Taking the expectation of this quantity over the stationary distribution of states, we obtain

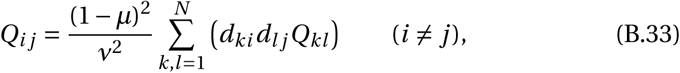

and *Q*_*ij*_ = 1 when *i* = *j*. Eq. (B.33) is valid for any regular graph; all the *Q*_*ij*_ terms can be found by solving a system of *N*(*N* − 1)/2 equations (since *Q*_*ij*_ = *Q*_*ji*_). We can also write eq. (B.5) in matrix form:

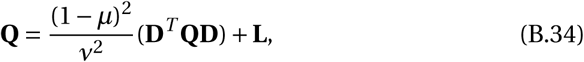

where **D** is the adjacency matrix of the dispersal graph (with elements *d*_*ij*_), ^*T*^ denotes transposition, and **L** is a diagonal matrix whose *i*th diagonal element is such that *Q*_*ii*_ = 1.

##### B.3.1 Transitive undirected graphs

When the dispersal graph is transitive, then all the elements on the diagonal of **L** are equal, so we can write 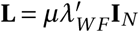, where **I**_*N*_ is the *N* by *N* identity matrix. Like in the case of a Moran updating, when the graph is also undirected, **D** = **D**^*T*^, and we also show by induction that **DQ** = **QD** (Grafen & Archetti, 2008).

Let us assume without loss of generality that initially (*t* = 0) all individuals are IBD (*q*_*ij*_(0) = **1**_*NN*_, where **1**_*NN*_ is the *N*-by-*N* matrix containing only ones) and of type *B* (*X*(0) = {0,…,0}). Also, let us denote by ζ_0_(*X*, *t*) the probability that the population is in state *X* at time *t* given that it was in state {0,…,0} at time 0, and by **E**_*t*_[] expectations with respect to that distribution, at time *t*. Then from eq. (B.32), since *q*_*ii*_ = 1, and given that the graph is regular,

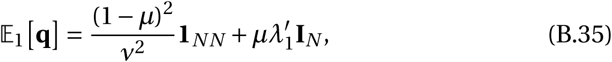

so

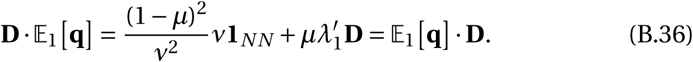

Then, assuming that **D** and **E**_t_[**q**] commute, and given that we assume an undirected dispersal graph (**D** = **D**^*T*^),

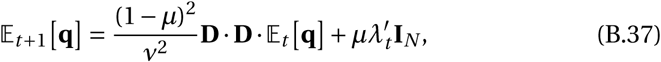

so

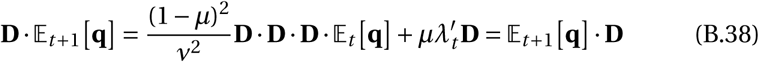

And so, when *t* → ∞, we have **D** ‧ **Q** = **Q** ‧ **D**.

Then with a transitive undirected dispersal graph, eq. (B.34), simplifies into

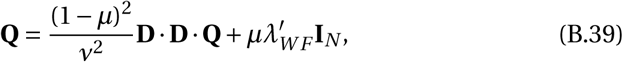

and so (for *μ* > 0),

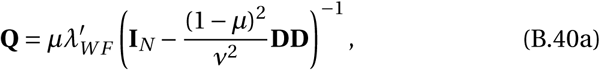

with

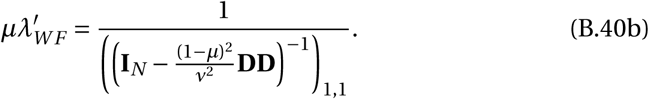

It is possible to find more explicit formulae when the graphs are transitive and when they are *n*-dimensional, and we do so for 1-D and 2-D graphs.

###### More about 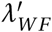

Like we did for the Moran model, we denote by **u** the *N* long column vector of ones. Using eq. (B.39), the fact that **D** and **Q** commute, and the regularity of the 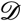 graph, we have

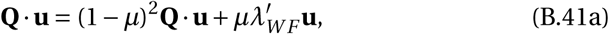

which simplifies into

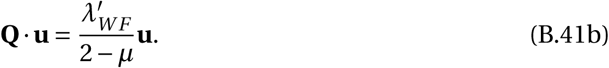

##### B.3.2 One-dimensional graphs

In a 1D graph, we can rewrite eq. (B.33) as follows, were 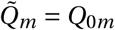 (numbering being done modulo *N*):

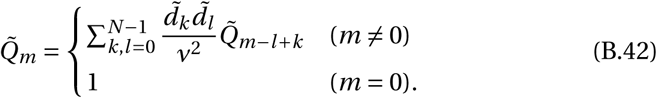

Using a Discrete Fourier Transform (see eq. (B.22)), we obtain,

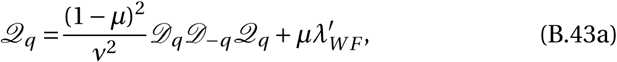

with

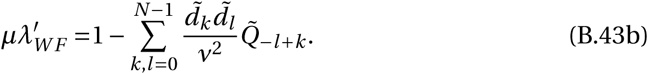

Solving for 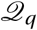, we obtain

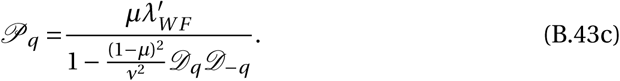

Then using an Inverse Fourier Transform to recover 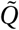 (see eq. (B.24)), we obtain

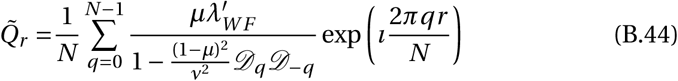

Noting that 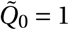, we can evaluate *λ*:

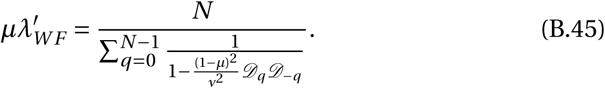

##### B.3.3 Two-dimensional graphs

Following the same method as previously, we obtain

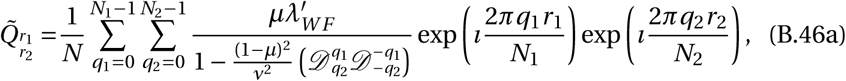

with

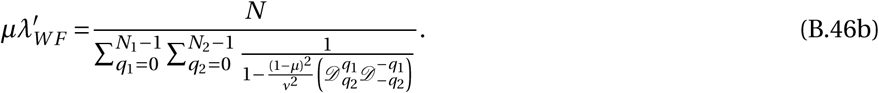

##### B.3.4 Illustration: Circle graph with self-loops

On a circle graph with self-loops (like in figure 3(b)), the Fourier transform of the dispersal distance is

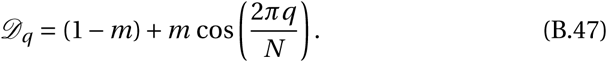

(Here *v* = 1, while with the circle graph we had *ν* = 2; this does not matter for IBD). We can evaluate 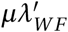 using eq. (B.45),

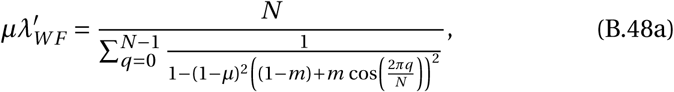

and when population size is infinite, this becomes

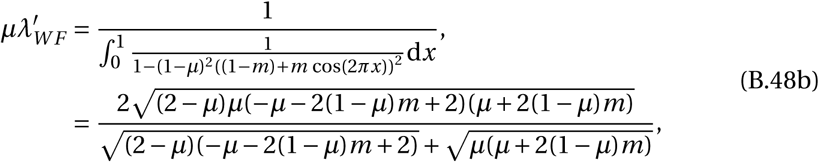

according to Mathematica (Wolfram Research, Inc., 2015). Here as well, the integral does not converge when *μ* → 0. Finally, we compute probabilities of identity by descent using eq. (B.46a), and obtain eq. (27) in the main text for neighbors on the the circle (*q* = 1).

## Appendix C

### C Specific life-cycles

#### C.1 Birth-Death updating

##### C.1.1 Derivatives of *B*_*ij*_ and *D*_*j*_

We will need to specify whether we consider two different sites, or the same site twice (*j* = *l*), and to this end introduce notation

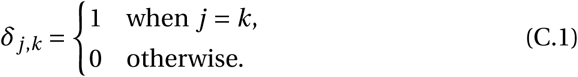

With a Moran Birth-Death updating rule (see eq. (14)), the derivatives of *B*_*ij*_ with respect to *f*_*l*_ is

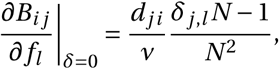

so that

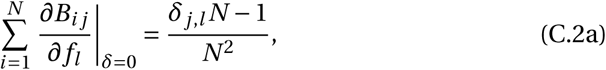

(because of the graph’s regularity, eq. (1)), and for *D*_*j*_ we obtain

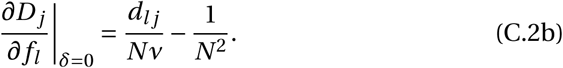

##### C.1.2 Expected frequency of social individuals

We replace the derivatives of *B*_*ij*_and *D*_*j*_ by their formulas for the Birth-Death life-cycle (eq. (C.2)), noting that for all *j*, *Q*_*jj*_ = 1 and remembering that *B*^*^ = 1/*N*; then eq. (10) becomes

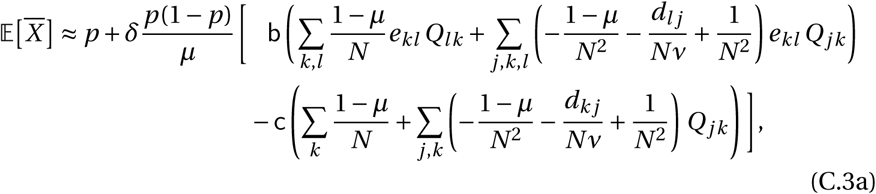

which after simplification becomes

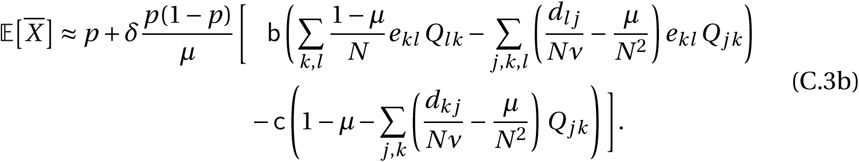

Using matrix notation, we obtain

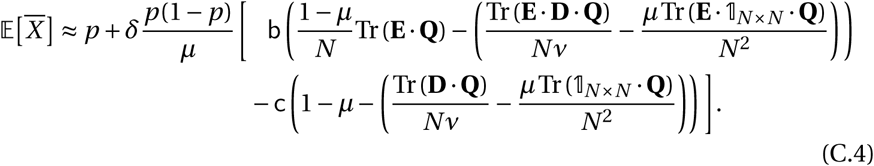

##### C.1.3 On transitive undirected dispersal graphs

We need results on **Q**, given by eq. (B.11) and eq. (B.14); they imply that

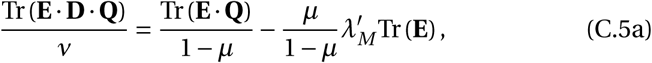

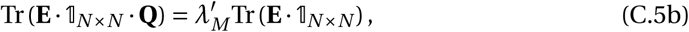

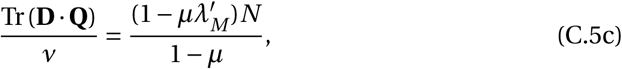

and

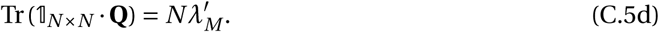

Back to eq. (C.4), we obtain

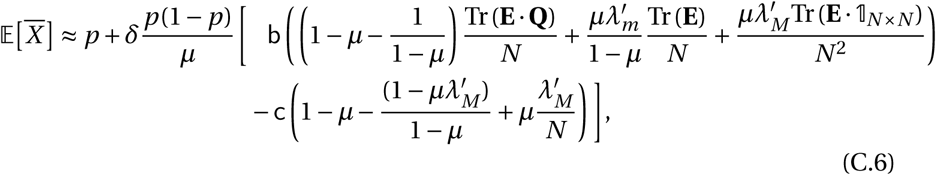

which simplifies into eq. (18) in the main text.

Now we show that the factor associated to b is negative when we exclude interactions with one-self (*i.e.*, when Tr(**E**) = 0). We use the following upper bounds: for all pairs of sites *i* and *j*, *Q*_*ij*_ ≤ 1, and *λ*′_*M*_ ≤ *N* (see eq. (B.19)). As a result, we have

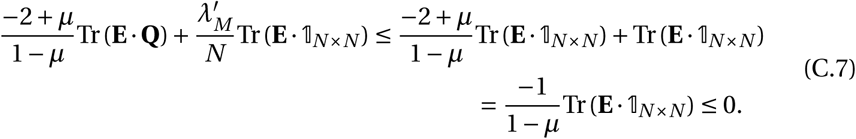

###### When the dispersal and interaction graphs are the same

We can further simplify eq. (18), using eq. (C.5d) again, and we obtain

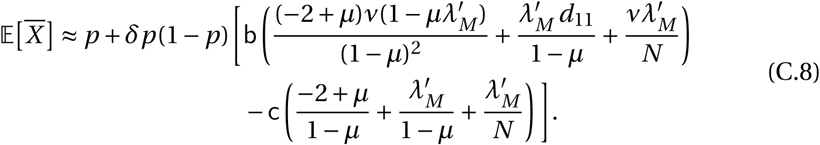

#### C.2 Death-Birth updating

##### C.2.1 Derivatives of *B*_*ij*_ and *D*_*j*_

With a Moran Death-Birth updating rule (see eq. (19)), the derivatives of *B*_*ij*_ and *D*_*j*_ with respect to *f*_*k*_ are given by the following equations:

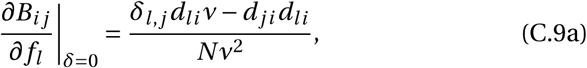

so that

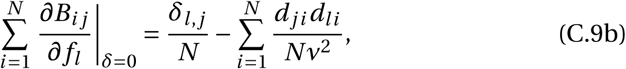

and

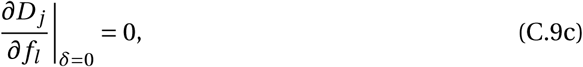

with *δ*_*k,j*_ as defined in eq. (C.1).

##### C.2.2 Expected frequency of social individuals

We replace the derivatives of *B*_*ij*_ and *D*_*j*_ by their formulas for the Death-Birth life-cycle (eq. (C.9)), noting that for all *j*, *Q*_*jj*_ = 1 and remembering that *B*^*^ = 1/*N*; then eq. (10) becomes

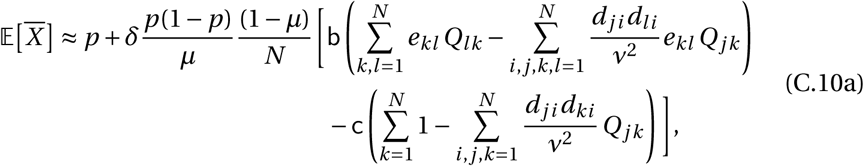

and under matrix form, we have

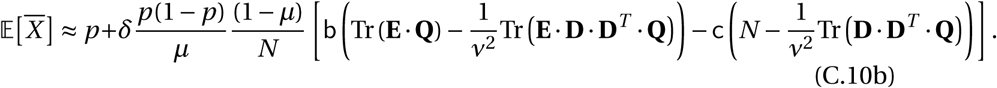

##### C.2.3 On transitive undirected dispersal graphs

We need results on **Q** given by eq. (B.13); it implies

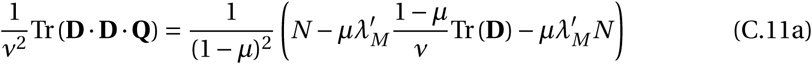

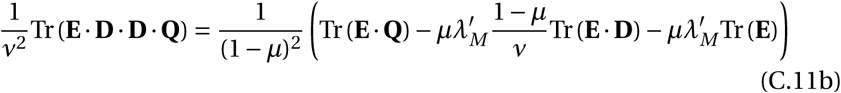

Plugging these into eq. (C.10b), we obtain eq. (21).

###### When the dispersal and interaction graphs are the same

We can further simplify eq. (21), using eq. (C.5d) again, and we obtain

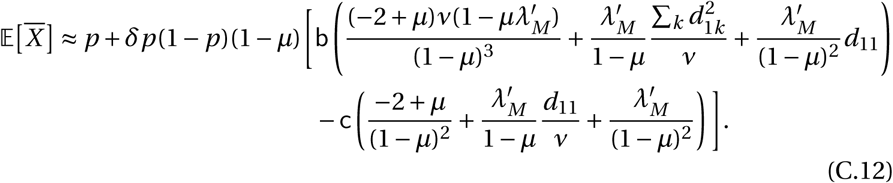

#### C.3 Wright-Fisher updating

##### C.3.1 Derivatives of *B*_*ij*_ and *D*_*j*_

With a Wright-Fisher updating rule (see eq. (22)), the derivatives of *B*_*ij*_ and *D*_*j*_ with respect to *f*_*k*_ are

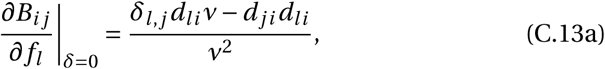

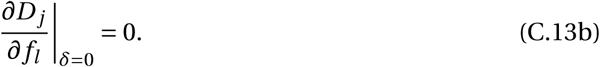

with *δ*_*k*,*j*_ as defined in eq. (C.1). This differs by a factor *N* from the Death-Birth version (eq. (C.9)), and since *B*^*^ = 1 (instead of 1/*N*) in the Wright-Fisher model, we end up with eq. (C.10b) for the expected frequency of type-*A* individuals in the population.

##### C.3.2 On transitive undirected dispersal graphs

Probabilities of identity by descent **Q** are not the same under a Wright-Fisher model as under a Moran model. Using the relationship between **Q** and **D** given in eq. (B.39), we can further simplify 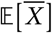 for this we need the following quantities:

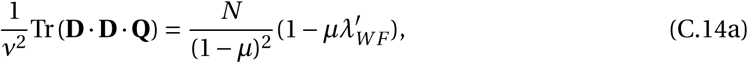

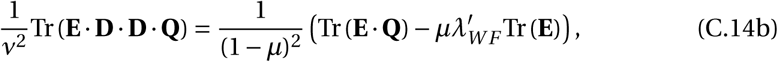

and so

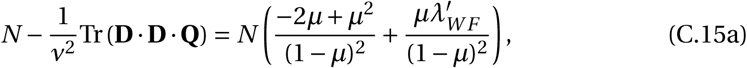

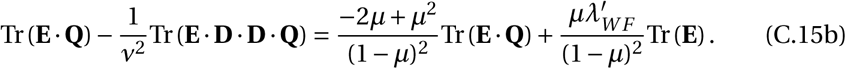

Plugging these into eq. (C.10b), we recover eq. (25).

---

**Figure S1:**
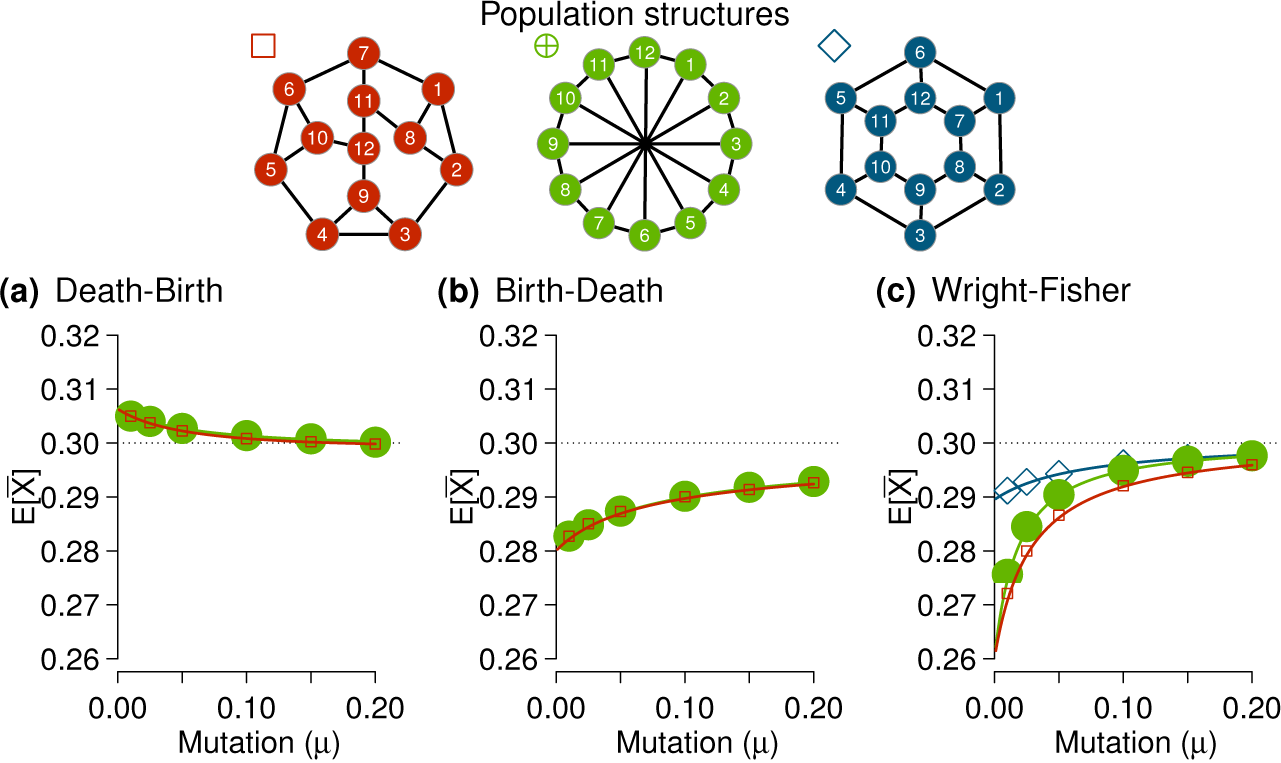
Expected frequency of type-A individuals 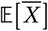, depending on population structure (legend on the first line), updating rule ((a): Moran Death-Birth, (b): Moran Birth-Death, (c): Wright-Fisher), and mutation probability *μ* (horizontal axis): Comparison between the theoretical prediction (curves) and the outcomes of numerical simulations (points). The horizontal dotted gray line corresponds to *p*, the expected frequency of type-*A* individuals when there is no selection (i.e., when *δ* = 0). Other parameters: *δ* = 0.005, *p* = 0.3, b = 8, c = 1.

## Footnotes

1 Here we extend the notation used in Allen et al. (2015), because in their study, *α*: *R* → {1, …, *N*}

## References

Allen, B. & Nowak, M. A. 2014: Games on graphs. EMS surveys in mathematical sciences 1(1):113–151.

Allen, B.; Nowak, M. A. & Dieckmann, U. 2013: Adaptive dynamics with interaction structure. The American Naturalist 181(6):E139–E163.

Allen, B.; Sample, C.; Dementieva, Y.; Medeiros, R. C.; Paoletti, C. & Nowak, M. A. 2015: The molecular clock of neutral evolution can be accelerated or slowed by asymmetric spatial structure. PLoS Comput Biol 11(2):1–32.

Allen, B.; Traulsen, A.; Tarnita, C. E. & Nowak, M. A. 2012: How mutation affects evolutionary games on graphs. Journal of Theoretical Biology 299:97–105. Evolution of Cooperation.

Caswell, H. 2001: Matrix population models. Wiley Online Library.

Champagnat, N.; Ferrière, R. & Méléard, S. 2006: Unifying evolutionary dynamics: from individual stochastic processes to macroscopic models. Theoretical population biology 69(3):297–321.

Champagnat, N. & Lambert, A. 2007: Evolution of discrete populations and the canonical diffusion of adaptive dynamics. The Annals of Applied Probability 17(1):102–155.

Cockerham, C. C. & Weir, B. S. 1993: Estimation of gene flow from F-statistics. Evolution 47(3):855–863.

Débarre, F.; Hauert, C. & Doebeli, M. 2014: Social evolution in structured populations. Nature Communications 5.

Frank, S. A. 1997: The price equation, fisher’s fundamental theorem, kin selection, and causal analysis. Evolution 51(6):1712–1729.

Gandon, S. & Rousset, F. 1999: Evolution of stepping-stone dispersal rates. Proceedings of the Royal Society of London B: Biological Sciences 266(1437):2507–2513.

Geritz, S.; Kisdi, E.; Meszena, G. & Metz, J. 1997: Evolutionarily singular strategies and the adaptive growth and branching of the evolutionary tree. Evolutionary Ecology 12(1):35–57.

Grafen, A. & Archetti, M. 2008: Natural selection of altruism in inelastic viscous homogeneous populations. Journal of Theoretical Biology 252(4):694–710.

Hindersin, L. & Traulsen, A. 2015: Most undirected random graphs are amplifiers of selection for birth-death dynamics, but suppressors of selection for death-birth dynamics. PLoS Comput Biol 11(11):1–14.

Kimura, M. & Crow, J. F. 1964: The number of alleles that can be maintained in a finite population. Genetics 49(4):725–738.

Lehmann, L. 2012: The stationary distribution of a continuously varying strategy in a class-structured population under mutation–selection–drift balance. Journal of Evolutionary Biology 25(4):770–787.

Lehmann, L.; Keller, L. & Sumpter, D. J. T. 2007: The evolution of helping and harming on graphs: the return of the inclusive fitness effect. Journal of Evolutionary Biology 20(6):2284–2295.

Lehmann, L. & Rousset, F. 2010: How life history and demography promote or inhibit the evolution of helping behaviours. Philosophical Transactions of the Royal Society B: Biological Sciences 365(1553):2599–2617.

Lehmann, L. & Rousset, F. 2014: The genetical theory of social behaviour. Philosophical Transactions of the Royal Society of London B: Biological Sciences 369(1642).

Lieberman, E.; Hauert, C. & Nowak, M. A. 2005: Evolutionary dynamics on graphs. Nature 433(7023):312–316.

Maciejewski, W. 2014: Reproductive value in graph-structured populations. Journal of Theoretical Biology 340:285–293.

Malécot, G. 1975: Heterozygosity and relationship in regularly subdivided populations. Theoretical Population Biology 8(2):212–241.

McAvoy, A. & Hauert, C. 2015: Structural symmetry in evolutionary games. Journal of The Royal Society Interface 12(111).

Moran, P. 1962: The statistical processes of evolutionary theory. Clarendon Press; Oxford University Press.

Nakamaru, M. & Iwasa, Y. 2006: The coevolution of altruism and punishment: Role of the selfish punisher. Journal of Theoretical Biology 240(3):475–488.

Nowak, M.; Sasaki, A.; Taylor, C. & Fudenberg, D. 2004: Emergence of cooperation and evolutionary stability in finite populations. Nature 428(6983):646–650.

Nowak, M. A. 2006: Five rules for the evolution of cooperation. Science 314(5805):1560–1563.

Nowak, M. A.; Tarnita, C. E. & Wilson, E. O. 2010: The evolution of eusociality. Nature 466(7310):1057–1062.

Ohtsuki, H.; Hauert, C.; Lieberman, E. & Nowak, M. A. 2006: A simple rule for the evolution of cooperation on graphs and social networks. Nature 441(7092):502–505.

Ohtsuki, H. & Nowak, M. A. 2006: The replicator equation on graphs. Journal of Theoretical Biology 243(1):86–97.

Ohtsuki, H.; Nowak, M. A. & Pacheco, J. M. 2007: Breaking the symmetry between interaction and replacement in evolutionary dynamics on graphs. Phys. Rev. Lett. 98:108106.

Queller, D. C. 1994: Genetic relatedness in viscous populations. Evolutionary Ecology 8:70–73. 10.1007/BF01237667.

Rousset, F. 2003: A minimal derivation of convergence stability measures. Journal of Theoretical Biology 221(4):665–668.

Rousset, F. 2004: Genetic Structure and Selection in Subdivided Populations. Princeton University Press, Princeton, NJ.

Rousset, F. & Billiard, S. 2000: A theoretical basis for measures of kin selection in subdivided populations: finite populations and localized dispersal. Journal of Evolutionary Biology 13(5):814–825.

Slatkin, M. 1991: Inbreeding coefficients and coalescence times. Genetical research 58(02):167–175.

Slatkin, M. 1993: Isolation by distance in equilibrium and non-equilibrium populations. Evolution 47(1):264–279.

Tarnita, C. E.; Ohtsuki, H.; Antal, T.; Fu, F. & Nowak, M. A. 2009: Strategy selection in structured populations. Journal of Theoretical Biology 259(3):570–581.

Tarnita, C. E. & Taylor, P. D. 2014: Measures of relative fitness of social behaviors in finite structured population models. The American Naturalist 184(4):477–488.

Taylor, P. 2010: Birth–death symmetry in the evolution of a social trait. Journal of Evolutionary Biology 23(12):2569–2578.

Taylor, P.; Lillicrap, T. & Cownden, D. 2011: Inclusive fitness analysis on mathematical groups. Evolution 65(3):849–859.

Taylor, P. & Maciejewski, W. 2012: An inclusive fitness analysis of synergistic interactions in structured populations. Proceedings of the Royal Society B: Biological Sciences.

Taylor, P. D. 1990: Allele-frequency change in a class-structured population. The American Naturalist 135(1):pp. 95–106.

Taylor, P. D. 1992: Inclusive fitness in a homogeneous environment. Proceedings of the Royal Society of London. Series B: Biological Sciences 249(1326):299–302.

Taylor, P. D.; Day, T. & Wild, G. 2007a: Evolution of cooperation in a finite homogeneous graph. Nature 447(7143):469–472.

Taylor, P. D.; Day, T. & Wild, G. 2007b: From inclusive fitness to fixation probability in homogeneous structured populations. Journal of Theoretical Biology 249(1):101–110.

Van Cleve, J. 2015: Social evolution and genetic interactions in the short and long term. Theoretical Population Biology 103:2–26.

Wakano, J. Y. & Lehmann, L. 2014: Evolutionary branching in deme-structured populations. Journal of theoretical biology 351:83–95.

Wild, G. & Traulsen, A. 2007: The different limits of weak selection and the evolutionary dynamics of finite populations. Journal of Theoretical Biology 247(2):382–390.

Wolfram Research, Inc. 2015: Mathematica Edition: Version 10.1. Wolfram Research, Inc, Champaign, Illinois.

